# PCB126-mediated effects on adipocyte energy metabolism and adipokine secretion may result in abnormal glucose uptake in muscle cells

**DOI:** 10.1101/2020.07.07.192245

**Authors:** Audrey Caron, Fozia Ahmed, Vian Peshdary, Léa Garneau, Ella Atlas, Céline Aguer

## Abstract

**Background:** Exposure to coplanar polychlorinated biphenyls (PCBs) is linked to the development of insulin resistance. Previous studies suggested that PCB126 alters muscle mitochondrial function through an indirect mechanism. Since PCBs are stored in fat, we hypothesized that PCB126 alters adipokine secretion, which in turn affects muscle metabolism.

**Objectives:** The objectives of this study were: 1) To study the impacts of PCB126 exposure on adipocyte cytokine/adipokine secretion; 2) To determine whether adipocyte-derived factors alter glucose metabolism and mitochondrial function in myotubes when exposed to PCB126; 3) To determine whether pre-established insulin resistance alters the metabolic responses of adipocytes exposed to PCB126 and the communication between adipocytes and myotubes.

**Method:** 3T3-L1 adipocytes were exposed to PCB126 (1-100 nM) in two insulin sensitivity conditions (insulin sensitive (IS) and insulin resistant (IR) adipocytes), followed by the measurement of secreted adipokines, mitochondrial function and insulin-stimulated glucose uptake. Communication between adipocytes and myotubes was reproduced by exposing C2C12 or mouse primary myotubes to conditioned medium (CM) derived from IS or IR 3T3-L1 adipocytes exposed to PCB126. Mitochondrial function and insulin-stimulated glucose uptake were then determined in myotubes.

**Results:** PCB126 significantly increased adipokine (adiponectin, IL-6, MCP-1, TNF-α) secretion and decreased mitochondrial function, glucose uptake and glycolysis in IR but not in IS 3T3-L1 adipocytes. Altered energy metabolism in IR 3T3-L1 adipocytes was linked to decreased phosphorylation of AMP-activated protein kinase (p-AMPK) and increased superoxide dismutase 2 levels, an enzyme involved in reactive oxygen species detoxification. Exposure of myotubes to CM from PCB126-treated IR adipocytes decreased glucose uptake, without altering glycolysis or mitochondrial function. Interestingly, p-AMPK levels were increased rather than decreased in myotubes exposed to the CM of PCB126-treated IR adipocytes.

**Conclusion:** Taken together, these data suggest that increased adipokine secretion from IR adipocytes exposed to PCB126 may explain impaired glucose uptake in myotubes.

## Introduction

Since 1990, the number of individuals with diabetes has quadrupled worldwide where in 2014, 422 million people were affected by this disease (Olokoba, Obateru, & Olokoba, 2012). Type 2 diabetes accounts for over 85% of diabetes cases (Diabètes Québec, Canadian Electronic Library, & Canadian Diabetes Association, 2011). This endocrine disorder is caused by insulin resistance in a number of tissues and organs including the liver, adipose tissue and skeletal muscles, followed by a dysfunction of pancreatic β-cells. The development of insulin resistance is multifactorial, involving the interaction of genes and environmental-behavioural risk factors.

Due to its mass, skeletal muscle is responsible for up to 80% of postprandial glucose disposal (Ferrannini et al., 1985) and it is therefore a key player in the development of insulin resistance and type 2 diabetes (Mogensen et al., 2007). Insulin resistance is also related to muscle mitochondrial dysfunction, which can be caused by oxidative stress (Bonnard et al., 2008). In addition, the accumulation of visceral fat is also recognized as a major cause in the development of insulin resistance and type 2 diabetes (Matsuda & Shimomura, 2013). Hypertrophied adipose tissue leads to increased inflammation and metabolic imbalance (Papers et al., 2005). Furthermore, alteration in the secretion of cytokines/adipokines by dysfunctional adipose tissue, such as tumor necrosis factor α (TNF-α), leptin and adiponectin, increases oxidative stress and induces mitochondrial dysfunction and insulin resistance in other tissues including skeletal muscle (Carey et al., 2006; Scherer, 2006; Steinberg, 2007). Therefore, altered adipose tissue-derived factors may affect normal adipose-to-muscle communication and therefore lead to the development of insulin resistance and type 2 diabetes.

It has been recently hypothesised that exposure to environmental pollutants may explain the sharp increase in the prevalence of type 2 diabetes. Epidemiological studies found a correlation between exposure to various persistent organic pollutants (POPs), such as polychlorinated biphenyls (PCBs) and p,p-dichlorodiphenylchloroethane (DDT), the development of insulin resistance and type 2 diabetes (Everett et al., 2007; Neel & Sargis, 2011; Sargis, 2014). PCBs are very stable compounds that accumulate in adipose tissue due to their lipophilicity. The phenyl rings and/or chlorine substituents arrangements dictate PCB mechanism of action. Dioxin-like PCBs (coplanar), such as PCB77 and PCB126, are associated with a wide range of toxic effects, such as reproductive dysfunction, immunotoxicity, liver damages, metabolic dysregulation, and developmental defects (WHO, 2010). Their toxicity is mainly mediated through the activation of aryl hydrocarbon receptor (AhR). Non-dioxin-like PCBs are also associated with liver damage, developmental and neurological effects, but their toxicity is not via AhR activity (Giesy & Kurunth, 2002). Exposure to PCBs most commonly found in the environment; PCB77, 118, 126 and 153, or mixtures of PCBs (e.g. Aroclor 1254); may be in part responsible for the current increased prevalence of metabolic dysfunctions and type 2 diabetes (Everett et al., 2007; Neel & Sargis, 2011; Rains & Jain, 2011; Ruzzin et al., 2010; Sargis, 2014). Exposure to Aroclor 1254, which contains multiple PCB congeners, induced hyperinsulinemia and insulin resistance in mice (Gray, Shaw, Gagne, & Chan, 2013). Furthermore, coplanar PCB exposure is linked to mitochondrial dysfunction, inflammation, and insulin resistance in mice, as well as in adipocytes and human umbilical vascular endothelial cells (Baker et al., 2013; Ruzzin et al., 2010; Wang, Lv, & Du, 2010). Moreover, diet-induced insulin resistance, adiposity, and adipose tissue inflammation in rats was exacerbated by pollutants (Gray et al., 2013; Lim et al., 2009).

Despite the growing body of evidence supporting the role of coplanar PCBs in the development of insulin resistance and type 2 diabetes, the effects of PCB exposure on the insulin sensitivity and energy metabolism of skeletal muscle has not been thoroughly investigated. Our group demonstrated that a 24hr-exposure of L6 skeletal muscle cells to PCB126 induced a 20% decrease in glucose uptake and glycolysis (Mauger, Nadeau, Caron, Chapados, & Aguer, 2016). This altered glucose metabolism in PCB126-treated L6 myotubes was consistent with another study showing decreased GLUT4 translocation and glucose uptake in muscle of rats exposed to a mixture of PCBs (Williams et al., 2013). Furthermore, we recently demonstrated that a single exposure of rats to PCB126 resulted in a reduced mitochondrial function measured in permeabilized muscle fibers, associated with an alteration in the expression levels of a number of enzymes involved in reactive oxygen species (ROS) detoxification (Tremblay-Laganière et al., 2019). However, no mitochondrial dysfunction was detected *in vitro* in L6 muscle cells exposed directly to PCB126 (Mauger et al., 2016). Skeletal muscle response to PCB126 is therefore different *in vivo* (whole organism) and *in vitro* (direct exposure of muscle cells to PCB126). Taken together, these studies suggest that the development of mitochondrial dysfunction in skeletal muscle of rats exposed to PCB126 was not the result of a direct effect of the pollutant on muscle mitochondria.

Since PCBs mostly accumulate in adipose tissue (Imbeault, Tremblay, Simoneau, & Joanisse, 2002), we hypothesized that PCB126 might first induce adipose tissue dysfunction resulting in altered adipokine and inflammatory cytokine secretion, which in turn could induce mitochondrial dysfunction in skeletal muscle. The overall aim of the present study was thus to study the role of adipose-to-muscle communication in PCB126-induced metabolic defects. The specific objectives were: 1) To determine the effect of PCB126 exposure on adipocyte cytokine/adipokine production; 2) To test whether the communication between adipose tissue and muscle explains a) abnormal muscle glucose metabolism and b) muscle mitochondrial dysfunction when exposed to PCB126; and 3) To determine whether insulin resistance in adipocytes alters their responses to PCB126 and alters the communication between adipocytes and muscle cells.

## Methods

### Cell culture

All cell lines were cultured in a humidified incubator (Thermo Scientific Forma Steri-cycle CO_2_ incubator) at 37°C with 5% CO_2_.

#### 3T3-L1 adipocytes

3T3-L1 (ATCC^®^ CL-173™) were grown in low glucose DMEM (1.0 g/L glucose, 4 mM L-glutamine and 110 mg/L sodium pyruvate, Wisent), 10% calf serum (CS, Wisent) and 1X antimycotic-antibiotic (AA) (Wisent). The medium was changed every two days until cells reached ~90% confluence. 3T3-L1 cells were then differentiated into adipocytes for 10 days using three different adipocyte differentiation media (ADM1, ADM2 and ADM3) at specific time as described in Table 1. Insulin resistance (IR) was induced by using differentiation media with high concentrations of insulin (500 nM, Sigma-Aldrich). For insulin sensitive (IS) adipocytes, the concentration of insulin in ADM1 and ADM2 was 100 nM and no insulin was present in ADM3 (Table 1).

**Table 1:**
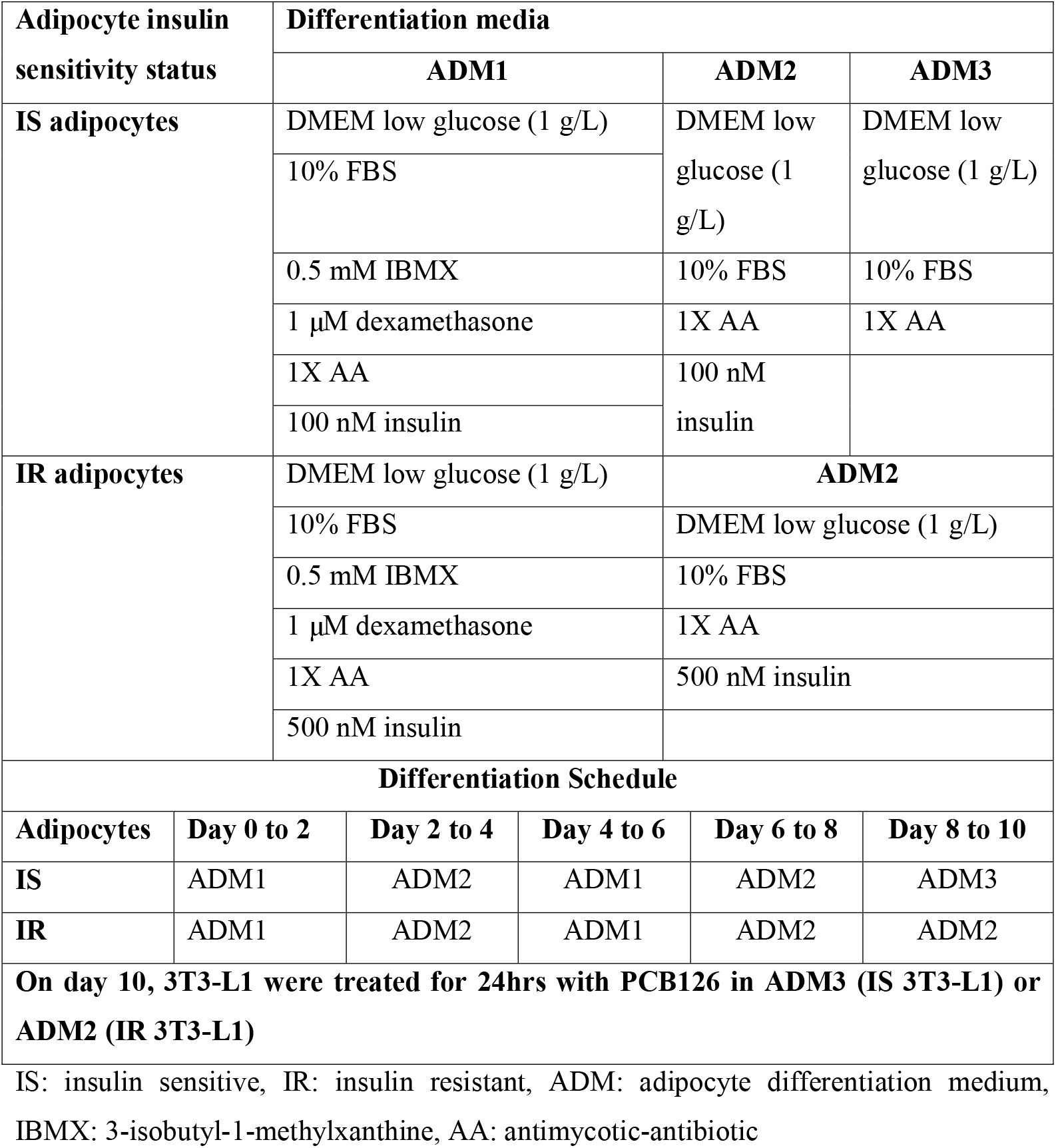
Adipocyte (3T3-L1) differentiation protocol and schedule.

#### C2C12 muscle cells

C2C12 myoblasts (Sigma-Aldrich) were grown in low glucose DMEM, 10% fetal bovine serum (FBS, Wisent) and 1X AA. The medium was refreshed every other day until cells reached ~90% confluence. C2C12 cells were then differentiated into myotubes for 7 days in DMEM low glucose, 2% FBS and 1X AA. Differentiation medium was refreshed every 2 or 3 days.

#### Mouse primary muscle cells

Mouse primary myoblasts derived from gastrocnemius and tibialis muscles of wild-type mice with a C57BL/6J background were a kind gift from Dr. Marc Foretz (Institut Cochin, Paris, France). Mouse primary muscle cells were cultured on matrigel (1X in DMEM, Corning) coated equipment in a DMEM:F12 1:1 medium (Wisent) supplemented with 20% FBS, 1X AA, 3 μg/mL gentamicin (Wisent) and 5 ng/mL recombinant mouse fibroblast growth factor basic (FGF-b, ThermoFisher). At ~90% confluence, mouse primary muscle cells were differentiated into myotubes for 7 days in low glucose DMEM supplemented with 2% FBS, 1X AA and 3 μg/mL gentamicin. Differentiation medium was refreshed every 2 or 3 days.

### PCB126 treatments

Treatments were done once the adipocytes were fully differentiated to study the impact of PCB126 on the secretome and metabolism of mature adipocytes. On day 10 of differentiation, adipocytes were exposed to 0, 1, 10 or 100 nM of PCB126 dissolved in 0.1% dimethyl sulfoxide (DMSO, Sigma-Aldrich) for 24hrs. These concentrations were chosen to represent environmentally relevant PCB126 concentrations. In fact, exposure to PCB126 in Canadian Inuit population is between 0.05 nM to 27 nM (Singh & Chan, 2017), while the daily intake of PCB126 is estimated to be 12 pg/day (USEPA, 2003). Control cells (0 nM PCB126) were exposed to 0.1% DMSO (vehicle). In order to determine whether some factors secreted by adipocytes upon exposure to PCB126 altered the metabolism of myotubes, after the 24hr-PCB126 treatment, the conditioned medium (CM) of adipocytes was used to treat differentiated C2C12 myotubes or mouse primary myotubes for 24hrs. This model was chosen over the co-culture model to study the unidirectional communication from adipocytes to myotubes. As a control condition, differentiated myotubes were also treated with the same concentrations of PCB126 in ADM2 (IR) or ADM3 (IS) for 24hrs (direct exposure). After the 24hr-treatments, cells were prepared for the different experiments described below.

### Cell viability

To determine whether the different treatments affected cell survival, viability was measured using the PrestoBlue method (ThermoFisher) according to manufacturer’s instructions. 3T3-L1 and C2C12 were grown and differentiated in 96-well plates (20,000 cells/well) and treated with PCB126 or 3T3-L1 CM for 24hrs as described above. Cells were then incubated for 30 min in 1X PrestoBlue^®^ reagent and absorbance was measured at 570 nm and 600 nm (reference wavelength) (BioTek instruments, Synergy™ HT Multi-Mode Microplate Reader). Each experiment was done in triplicate for 3T3-L1 adipocytes and in 6 replicates for C2C12 myotubes, and repeated on 3 independent cultures (n=3).

### Lipid accumulation

To determine whether the treatments had an effect on 3T3-L1 and C2C12 intracellular lipid accumulation, lipid droplets were stained by Oil Red O (Sigma-Aldrich). Lipid accumulation is an indicator of adipocyte differentiation. In muscle cells, intramyocellular lipid accumulation is believed to be related to insulin resistance development (Aguer et al., 2010; Krssak et al., 1999; Perseghin et al., 1999). For these experiments, 3T3-L1 were grown and differentiated in 6-well plates, while C2C12 were grown and differentiated in 12-well plates. The cells were treated with PCB126 and/or 3T3-L1 CM as described above, and then fixed with 4% paraformaldehyde (Alfa Aesar) in phosphate-buffered saline (PBS) for 10 min. Lipid droplets were stained with 0.5% Oil Red O dissolved in chloroform:ethanol (1:1, Fisher Scientific) for 10 min, as previously described (Aguer et al., 2010). Cells were washed three times with distilled water and visualized by light microscope (EVOS XL Core Imaging System) using a 40X objective and a constant light intensity. Oil Red O was then extracted with 70% isopropanol (Fisher Scientific) and absorbance read at 490 nm (Aguer et al., 2010). Each experiment was done in duplicate and repeated on 3 independent cell cultures (n=3).

### mRNA quantification by RT-qPCR

3T3-L1 were grown and differentiated in 6-well plates followed by 24hr-treatment with PCB126 as described above. Cells were then lysed using RNease lysis buffer (RLT buffer) with 1% mercaptoethanol (Sigma-Aldrich) from the RNeasy Mini Kit (Qiagen) and lysates were stored at −80°C. Total RNA was extracted from cell lysates using the RNase Mini Kit where genomic DNA was removed using the RNase-Free DNase Kit (Qiagen) following the manufacturer’s recommendations. RNA concentration and extraction quality were measured using a NanoDropTM 1000 Spectrophotometer (ThermoFisher). Then, 0.5 μg RNA was reverse transcribed with iScript cDNA Synthesis Kit (Bio-Rad) following the manufacturer’s protocol in a CFX96-PCR Detection System (Bio-Rad). Complementary DNA (cDNA) was amplified and quantified in a CFX96-PCR Detection System using the iQSYBR SsoFast EvaGreen Supermix (Bio-Rad). Primers for each target genes are summarized in Table 2. All genes were normalized to β-actin levels and analyzed using the comparative CT (ΔΔCT) method, as previously described (Schmittgen & Livak, 2008). Each independent experiment (n=3) was done in triplicate.

**Table 2:**
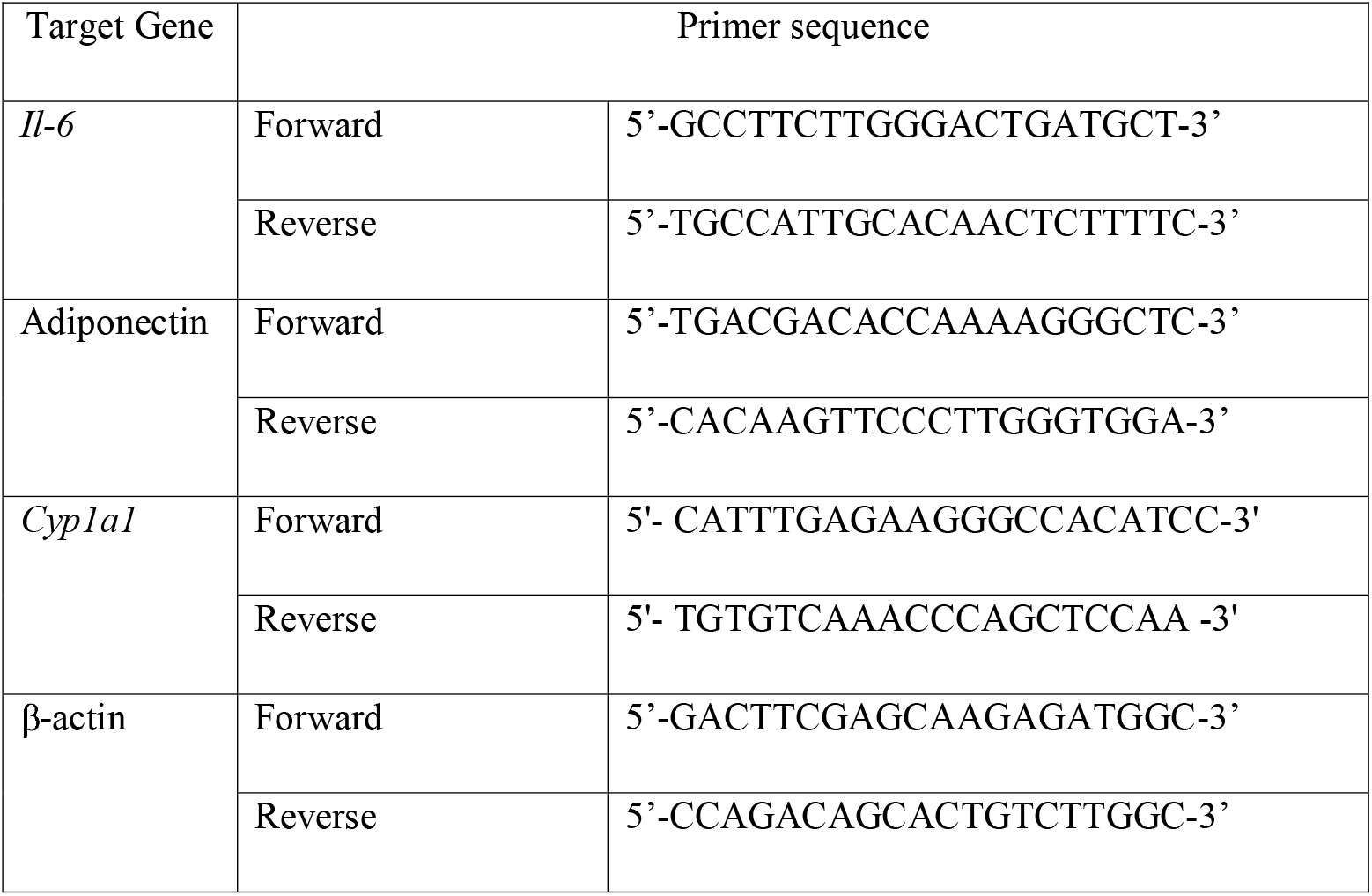
Primer sequence used for RT-qPCR.

### Adipokine measurements

To determine whether the different treatments affected cytokine and adipokine secretion, 3T3-L1 were grown and differentiated in 6-well plates followed by 24hr-treatments as described above. At the end of the treatments, culture media were collected and stored at −80°C until further cytokine/adipokine measurements. Cells were lysed in 0.05 M NaOH (Fisher Scientific) and protein quantified using the Bradford method (Bio-Rad). Cytokine/adipokine (IL-6, TNF-α, leptin, adiponectin, MCP-1, PAI-1, and resistin) concentrations in the different culture media were determined by the Bio-Plex method using the mouse adipocyte magnetic bead panel kit (Millipore, MADCYMAG-72K-07), according to manufacturer’s instructions (Millipore, 2013). Samples were diluted 20X in DMEM low glucose and 10 μL of each sample and standards were added to the detection beads and stirred rapidly at room temperature for 2hrs, washed 3 times followed by addition of secondary antibodies coupled to biotin for 30 minutes. After 3 washes, SA-PE (streptavidin-Phycoerythrin conjugate) was added to each well for 10 minutes at room temperature, rinsed 3 times and read by the Bio-Plex (Bio-Rad, Bio-Plex MAGPIX Multiplex Reader). Each experiment was done in duplicate and repeated with media from 3 independent experiments (n=3).

Adipokines measured with the Bioplex method are expressed relative to the vehicle condition since experiments were done independently for IS and IR conditions (preparation of samples and Bioplex experiments done at different times). In order to compare IS and IR conditions, and to confirm the Bioplex results, some adipokines were also re-measured, using ELISA kits, in the CM of adipocytes prepared in parallel for IS and IR conditions. Levels of adiponectin in the CM were determined after a 1:1000 dilution using the adiponectin mouse ELISA kit (Biomatik, EKL54022). Levels of IL-6 were measured after a 1:2 CM dilution using the IL-6 mouse ELISA kit (Thermo-Fisher, BMS603-2). Levels of leptin and TNF-α were determined in undiluted samples using the Leptin mouse ELISA kit (Biomatik, EKB01861) and the TNF-α mouse ELISA kit (Biomatik, EKA51917), respectively. Each measurement was done in duplicate and repeated with media from 3 or 4 independent experiments (n=3-4).

### Lipolysis

To determine whether PCB126 or insulin sensitivity status in adipocytes had an impact on lipolysis, glycerol and free fatty acids (FFA) were quantified in the adipocyte media by using a lipolysis quantification kit (ZenBIO, LIP-3-NC-L1) following the manufacturer’s protocol. Briefly, 3T3-L1 adipocytes were differentiated and treated with PCB126 as described above, in 96-well plates (20,000 cells/well). Then, they were incubated for 3hrs in assay buffer with PCB126. The media was used to quantify FFA and glycerol secretion. Absorbance was read at 540 nm. Each experiment was done in triplicate and repeated with media from 3 independent experiments (n=3).

### Mitochondrial respiration and glycolysis

3T3-L1 and C2C12 energy metabolism, mitochondrial function and glycolysis, were measured by determination of oxygen consumption rates (OCR) and of extracellular acidification rates (ECAR), respectively, using an extracellular flux analyzer (Seahorse XF-96, Agilent). The protocol provided by Seahorse Bioscience was followed for these experiments with slight modifications. At the end of the 24hr-treatments, cells were rinsed three times with assay buffer (8.3 g/L DMEM, 2 mM sodium pyruvate 5 mM dextrose, and 0.75 mM L-glutamine, pH 7.4, all from Sigma-Aldrich). Cells were then incubated at 37°C, without CO_2_, in 180 μL of assay buffer for 45 minutes. OCR and ECAR were first measured at baseline for 4 cycles including: 2 min-measurement, medium mixing for 2 min and 2 min-pause before starting the next cycle. Then, inhibitors of the respiratory chain were injected into each well in the following order: oligomycin, carbonyl cyanide-4-(trifluoromethoxy)phenylhydrazone (FCCP), and antimycin A (all from Sigma-Aldrich). After each injection, OCR and ECAR were measured for 3 cycles (measuring, mixing, rest; 2 min each). Final inhibitor concentrations used were 600 ng/mL oligomycin, 1 μM FCCP and 2 μM antimycin A for 3T3-L1 and 600 ng/mL oligomycin, 1 μM FCCP and 4 μM antimycin for C2C12. At the end of the experiment, cells were lysed in 50 μL of 0.05 M NaOH and proteins were quantified by the Bradford method. Mitochondrial OCR (non-mitochondrial OCR (antimycin A condition) subtracted from total OCR) and ECAR values are expressed per μg of total cellular protein. Each experiment was done in 5 or 6 replicates and repeated on 4 independent cultures (n=4).

### Glucose uptake

3T3-L1, C2C12 and mouse primary muscle cells were grown and differentiated in 24-well plates (100,000 cells/well) or 48-well plates (50,000 cells/well), followed by 24hr-treatments as described above. Glucose uptake was measured as in (Klip, Li, & Logan, 1984). After PCB126 treatment, cells were starved for 3hrs in serum-free DMEM. During the last 20 min of the starvation period, 100 nM insulin was added in half of the wells. Cells were then washed and incubated in Hepes Buffered Saline (HBS, 140 mM NaCl (Fisher Scientific), 20 mM Hepes-Na (Sigma-Aldrich), 5 mM KCl (Fisher Scientific), 2.5 mM MgSO_4_, (Fisher Scientific) and 1 mM CaCl_2_ (Sigma-Aldrich), pH 7.4) with 10 μM 2-deoxy-Glucose and 0.5 μCi/mL ^3^H-2-deoxy-glucose (Perkin Elmer). Cytocholasin B (5 μM, Sigma-Aldrich) was used to determine non-specific glucose uptake. Cells were lysed in 0.5 mL of 0.05 M NaOH and 0.4 mL were measured by scintillation counting with a Tri-Carb2910TR counter (Perkin Elmer). The remaining cell lysate was used to determine protein content using a Bradford protein assay. Each experiment was done in triplicate and repeated on 3 or 4 independent cultures (n=3-4).

### Western blots

3T3-L1 adipocytes and C2C12 myotubes were lysed in lysis buffer (20 mM Tris-HCl, 50 nM NaCl, 250 mM sucrose, 1% 100X triton, 50 mM NaF, 5 mM NaPP, 1 mM Na-orthovanadate). Twenty to 30 μg of proteins were separated by SDS-PAGE and then transferred to nitrocellulose membranes (GE Healthcare). Monoclonal anti-glutathione peroxidase 1 (GPx1, ab108427), monoclonal anti-glutathione peroxidase 4 (GPx4, ab125066), polyclonal anti-glutaredoxin 2 (Grx2, ab191292), MitoProfile^®^ Total OXPHOS Rodent WB Antibody Cocktail (anti-ATP5a, anti-complex III, anti-complex II) (MS604) (all from Abcam), monoclonal anti-catalase (D5N7V, #14097), polyclonal anti-α-tubulin (#2144), monoclonal glyceraldehyde-3-phosphate dehydrogenase (GAPDH, 14C10), monoclonal anti-AS160 (Akt substrate of 160 kDa, C6947, #2670S), polyclonal anti-phospho-AS160 (Thr 642, #4288S), polyclonal anti-phospho-IRS1 (insulin receptor substrate 1, S307, #2381S), polyclonal anti-phospho-Akt (S473, #9271S), monoclonal anti-Akt (C67E7, #4691S), polyclonal anti-phospho-GSK3α/β (glycogen synthase kinase 3, S2119, #9331S), polyclonal anti-phospho-AMPK (AMP activated protein kinase, #2535S), polyclonal anti-AMPK (#2532S) (all from Cell Signaling Technology), and polyclonal anti-superoxide dismutase (SOD2) (sc-30080, Santa Cruz), were used as primary antibodies at a dilution of 1:1000. The secondary antibodies used were anti-mouse (sc-516102) and anti-rabbit (sc-2357) antibodies coupled to horseradish peroxidase (Santa Cruz), diluted 1:5000. Proteins were visualized using SuperSignal West Pico Western Blot Kit (34580, Thermo Scientific) or Clarity Western ECL Substrate (170-5061, Biorad) and imaged using ChemiDoc™ Imager and VisionWorks LS (UVP). Expression of proteins was quantified by densitometry analysis using ImageJ program (National Institutes of Health, USA).

### Statistical analysis

Data shown are the means ± standard error of the mean (SEM) of at least 3 independent experiments. First, the average of the technical replicates within each independent experiment was determined, and then the mean of the 3 to 4 independent experiments were calculated. The SEM presented on the figures represents the SEM of the independent experiments. A Student’s t-test or one- or two-way ANOVA with Fisher’s protected least significant difference (PLSD) post-hoc test were used to determine statistical differences using Statview 5.0 Software (SAS Institute, USA) and GraphPad Prism version 6.0e (La Jolla, USA). With the Fisher’s PLSD post-hoc test used, we only compared PCB126-treated cells to the vehicle (0 nM PCB, 0.1% DMSO). A *P*<0.05 was considered significant.

## Results

### Effect of insulin concentrations on insulin sensitivity and lipid accumulation in differentiating 3T3-L1 adipocytes

In order to induce insulin resistance in 3T3-L1 adipocytes, 3T3-L1 cells were exposed to high concentrations of insulin (500 nM) during differentiation (IR condition). The IR condition significantly reduced insulin-stimulated glucose uptake in 3T3-L1 adipocytes compared to the IS condition (Figure S1A, fold increase in response to insulin: *P*=0.0438 IS vs IR). Moreover, there was no difference in lipid accumulation between IS and IR 3T3-L1 adipocytes, suggesting that the two differentiation protocols did not affect adipogenesis (Figure S1B).

### Effect of direct and indirect PCB126 exposure on lipid accumulation and cell viability in IS and IR adipocytes and C2C12 myotubes

We determined the effect of PCB126 or CM treatments on neutral lipid accumulation and cell viability in IS and IR 3T3-L1 adipocytes and C2C12 myotubes. As shown in Figure S2, neither PCB126/CM treatments, nor IS or IR conditions significantly altered intracellular lipid accumulation (Figure S2A) and cell viability (Figure S2B) in 3T3-L1 adipocytes and C2C12 myotubes.

### Effect of direct and indirect PCB126 exposure on *Cyp1a1* mRNA expression in IS and IR adipocytes and C2C12 myotubes

Coplanar PCBs are known to activate AhR, which in turn induces the transcription of cytochrome P450 1A (CYP1A) gene subfamily, leading, amongst other things, to increased inflammation (Baker et al., 2015; Wang et al., 2010). PCB126 exposure of 3T3-L1 adipocytes, at 100 nM for 24hrs, induced a 35-fold or a 85-fold increase in the expression of *Cyp1a1* mRNA in IS and IR conditions, respectively (Figure 1A, *P*=0.0026). No significant differences in *Cyp1a1* expression levels were observed in the IS versus IR conditions (Figure 1A). Direct exposure of C2C12 to PCB126 or exposure of C2C12 to CM from PCB126-treated adipocytes at the highest concentrations (10 and 100 nM) also increased *Cyp1a1* mRNA but this increase did not reach statistical significance (Figure S3).

**Figure 1.**
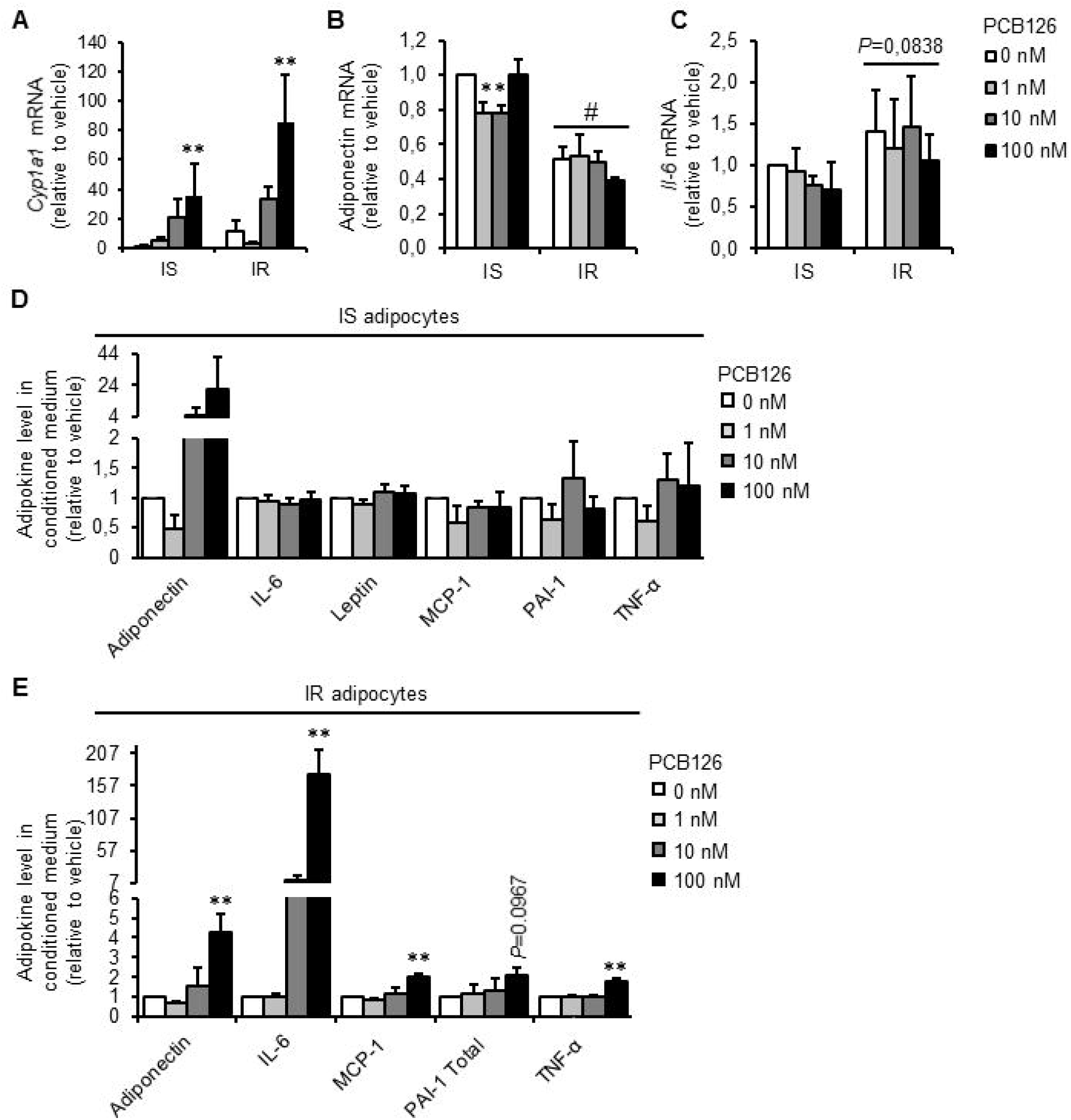
Effect of PCB126 exposure and insulin sensitivity conditions on adipokine expression and secretion in 3T3-L1 adipocytes. 3T3-L1 adipocytes were differentiated in insulin sensitive (IS) and insulin resistant (IR) conditions and treated for the last 24hrs of differentiation with different PCB126 concentrations. A-C. *Cyp1a1* (A), adiponectin (B), and *Il-6* (C) mRNA levels normalized to β-actin mRNA levels and analyzed using the ΔΔCT method. Average of normalized ΔΔCT is presented relative to the vehicle (IS, no PCB) ±SEM. (n=3 independent experiments, each independent experiment was done at least in triplicate, *: *P*<0.05 compared to 0 nM PCB126, #: *P*<0.05 compared to IS adipocytes). D-E. Adipokines secreted in the medium of 3T3-L1 adipocytes differentiated in IS (D) or IR (E) conditions. Data are presented relative to the vehicle as mean ±SEM. n=3 independent experiments, each independent experiment was done in 2 replicates. **: *P*<0.01 compared to 0 nM PCB126.

### Adipocytokine expression and secretion by adipocytes in response to PCB126 treatment in IS and IR conditions

We then determined whether a 24hr-PCB126 exposure altered the cytokine profile in IS and IR 3T3-L1 adipocytes by assessing mRNA expression and secretion of cytokines/adipokines. In IS adipocytes, 1 and 10 nM PCB126 exposure resulted in a significant decrease in adiponectin mRNA expression compared to the vehicle (*P*=0.0492 and 0.0397 respectively, Figure 1B). In IR adipocytes, PCB126 treatment did not significantly alter adiponectin mRNA expression, but there was a significant decrease in adiponectin mRNA expression in IR adipocytes compared to IS adipocytes, independently of PCB126 treatment (IR effect: *P*=0.0003, Figure 1B). *Il-6* mRNA expression was not altered by PCB126 exposure in both IS and IR adipocytes (Figure 1C), although in IR adipocytes *Il-6* mRNA expression trended to be increased compared to IS adipocytes but it did not reach statistical significance (IR effect: *P*=0.0838, Figure 1C). In addition, in IS adipocytes, PCB126 treatment did not significantly alter the secretion of cytokines/adipokines (Figure 1D). However, in IR adipocytes, a 100 nM PCB126 treatment significantly increased the secretion of adiponectin (*P*=0.0098), IL-6 (*P*=0.0002), MCP-1 (*P*=0.0018), and TNF-α (*P*<0.0001), and tended to increase the secretion of PAI-1 (*P*=0.0967) (Figure 1E). Leptin was not detected in the CM of IR adipocytes treated with 0, 1, and 10 nM PCB126, while it was when adipocytes were treated with 100 nM PCB126, suggesting an increased leptin secretion when exposed to 100 nM PCB126 (data not shown). Resistin was also measured in the CM of IS and IR adipocytes, but was not detected with the kit used. In this first set of experiments, IS and IR adipocytes were not prepared at the same time, preventing the comparison between the two differentiation conditions. In order to compare adipokine secretion between IS and IR conditions, and to confirm results obtained with the Bioplex method, some adipokines were also measured by ELISA kits. IL-6 levels were significantly higher in the medium of IR adipocytes compared to IS adipocytes, independently of PCB126 treatments (Figure S4A, IR effect: *P*=0.0062). Exposure to 100 nM PCB126 increased IL-6 secretion in both IS and IR adipocytes (PCB126 effect: *P*=0.0313), but the magnitude of the increase was much higher in IR adipocytes. Adiponectin levels were higher in the medium of IR compared to IS adipocytes, independently of PCB126 treatments (IR effect: *P*<0.001), whereas 10 nM PCB126 exposure significantly increased adiponectin levels in the medium of both IS and IR adipocytes (PCB126 effect: *P*=0.0093), but again, with a larger increase in IR compared to IS conditions (Figure S4B). Leptin levels were not significantly different between IS and IR conditions, and PCB126 treatments did not alter leptin levels in the medium of either IS or IR adipocytes (Figure S4C). This result does not confirm the increased leptin secretion in the medium of IR adipocytes measured with the Bioplex method. It is however important to note that leptin levels measured in the medium of adipocytes was very low and close to the detection level of the ELISA kit used, which can explain the absence of measured effects. TNF-α was also measured in the medium of adipocytes but was not detected with the ELISA kit used (data not shown).

### Effect of PCB126 treatment and insulin resistance on lipolysis

Another player in adipose-to-muscle communication is an altered secretion of fatty acids by adipose tissue (Rachek, 2014), which might be promoted by PCB exposure (Fræch et al., 2012; Regnier & Sargis, 2014) and/or insulin resistance. We therefore determined whether a 24hr-PCB126 exposure or insulin resistant conditions altered the rate of lipolysis in 3T3-L1 adipocytes. In IS adipocytes, exposure to 10 and 100 nM PCB126 for 24hrs decreased lipolytic rate compared to the vehicle condition (for [FFA]: *P*=0.049 and *P*=0.0147 for 10 and 100 nM PCB126, respectively) (Figure 2). However, in IR adipocytes, a 24hr-PCB126 treatment did not significantly alter lipolytic rate (Figure 2). In addition, there was a significant decreased lipolytic rate in IR adipocytes compared to IS adipocytes (IR effect: *P*=0.0009 and *P*=0.0005 for [FFA] and [glycerol], respectively) (Figure 2). This lower lipolytic rate in IR adipocytes was probably the result of the high insulin concentrations used to induce insulin resistance, since insulin is known to inhibit lipolysis and favor triglyceride synthesis (Chakrabarti et al., 2013; Cornelius, MacDouglad, & Lane, 1994).

**Figure 2.**
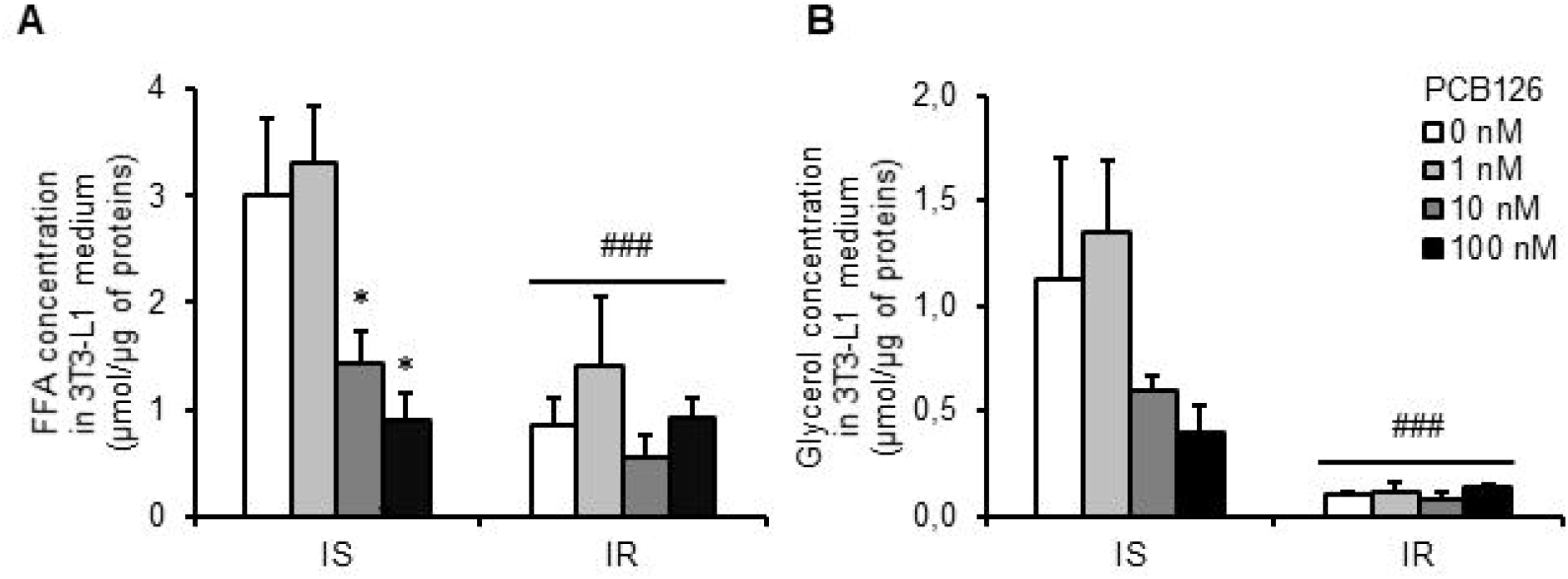
Effect of PCB126 exposure and insulin sensitivity conditions on lipolysis in 3T3-L1 adipocytes. 3T3-L1 adipocytes were differentiated in insulin sensitive (IS) and insulin resistant (IR) conditions and treated for the last 24hrs of differentiation with different PCB126 concentrations. As markers of lipolysis, free fatty acid (FFA) (A) and glycerol (B) levels were measured in the conditioned medium as described in the method section. Data are presented as mean ±SEM. n=3 independent experiments, each independent experiment was done in 2 replicates. *: *P*<0.05 compared to 0 nM PCB126 and ###: *P*<0.001 main effect of IR conditions.

### Effect of direct or indirect PCB126 exposure on mitochondrial function in IS and IR adipocytes and myotubes

#### Mitochondrial function in IS and IR adipocytes directly treated with PCB126

We first determined the effect of a 24hr-exposure to PCB126 in IS and IR conditions on adipocyte mitochondrial function. In IS adipocytes, OCR was not altered by a 24hr-PCB126 exposure (Figure 3A, left panel), while in IR adipocytes, 1 and 10 nM PCB126 resulted in a significant decrease in resting OCR and proton leak (state 4 OCR) (Figure 3A, right panel) (for 1 nM *P*=0.0279 and 0.0429 and for 10 nM *P*=0.0480 and 0.0448 for resting and proton leak OCR, respectively). To determine if this altered mitochondrial function in IR adipocytes exposed to PCB126 was the result of decreased levels of mitochondrial complexes, we measured the impact of PCB126 on the expression of the respiratory chain complexes (Figure 3B). Treatment with 100 nM PCB126 significantly increased ATP5A expression in IR adipocytes compared to vehicle (*P*=0.0065). A slight increase was also seen at 10 nM (*P*=0.0607). There was also a tendency for increased expression of complexes II and III with 24hr PCB126 exposure in IR adipocyte without reaching significance. Therefore, the decreased OCR in IR adipocytes exposed to PCB126 cannot be explained by a decrease in the levels of mitochondrial complexes. Rather, our results suggest that upon PCB126 exposure, IR adipocytes try to compensate the decreased mitochondrial function by increasing mitochondrial complex levels.

**Figure 3.**
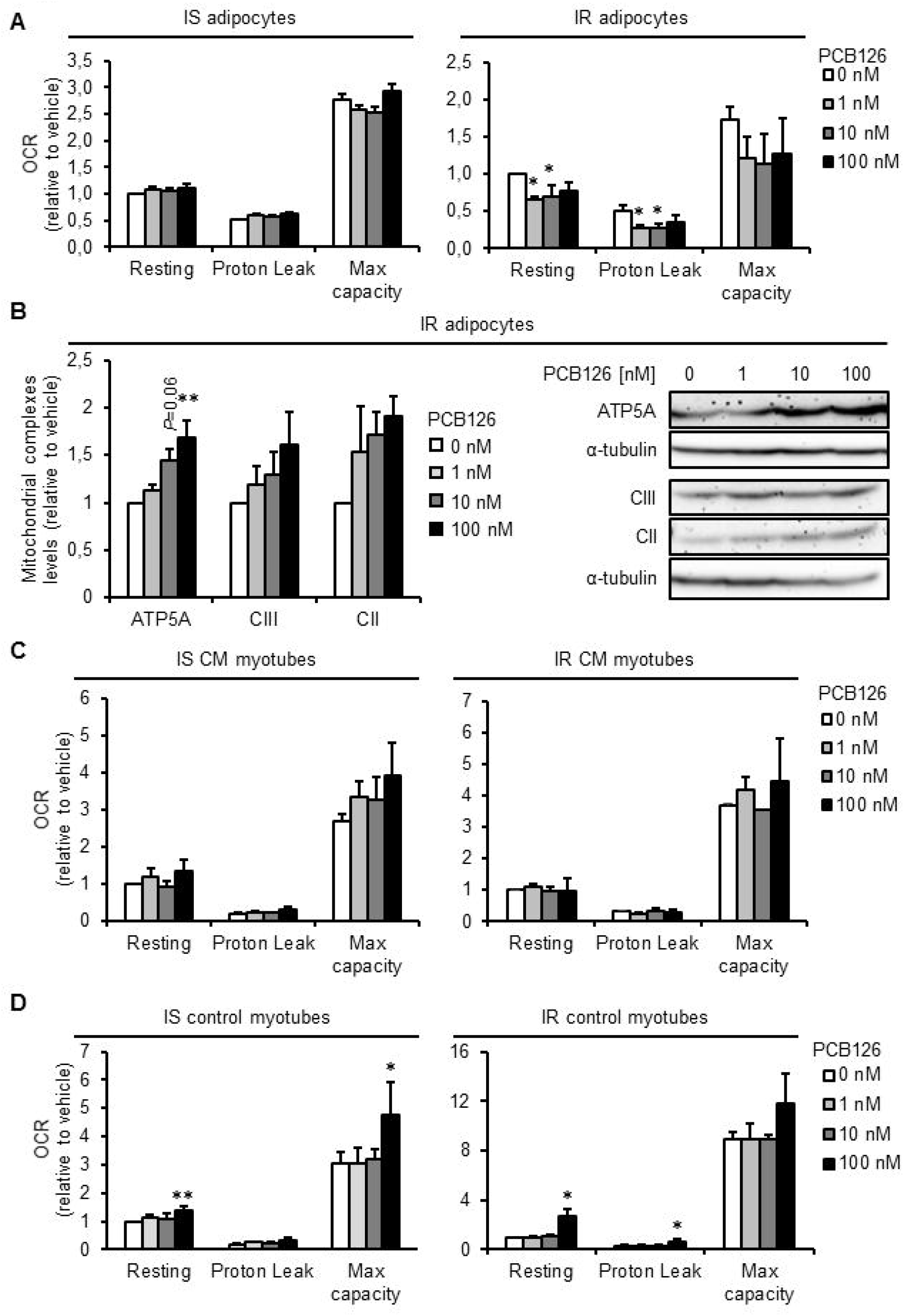
Mitochondrial function in 3T3-L1 adipocytes exposed to PCB126 in different insulin sensitivity conditions and in C2C12 myotubes exposed to the conditioned medium (CM) of insulin sensitive (IS) and insulin resistant (IR) adipocytes treated with PCB126. A. 3T3-L1 adipocytes were differentiated in IS (left panel) and IR (right panel) conditions and treated for the last 24hrs of differentiation with different PCB126 concentrations. B. Levels of mitochondrial complexes (complexes II and III and ATPase) in 3T3-L1 adipocytes differentiated in IR conditions and exposed for 24hrs to different concentrations of PCB126. Right panel: quantification by density analysis, left panel: representative western blots. α-tubulin was used a loading control. n=3 independent experiments. C. Differentiated C2C12 myotubes were exposed for the last 24hrs of differentiation to the CM of 3T3-L1 adipocytes exposed to different PCB126 concentrations in IS (left panel) or IR conditions (right panel). D. Differentiated C2C12 myotubes were directly exposed for the last 24hrs of differentiation to different PCB126 concentrations in IS (left panel) or IR conditions (right panel). A, C, and D. Oxygen consumption rates (OCR) were measured with a Seahorse analyzer (Agilent). OCR were first measured in resting conditions, and cells were treated subsequently with 600 ng/mL oligomycin, 1 μM carbonyl cyanide-4-(trifluoromethoxy)phenylhydrazone (FCCP), and 2 μM (for 3T3-L1) or 4 μM antimycin A (for C2C12) to determine OCR due to proton leak, maximal, and non-mitochondrial respiration, respectively. n=4 independent experiments, each independent experiment was done in 5 replicates. A-D. Data are presented relative to the vehicle as mean ±SEM. *: *P*<0.05, **: *P*<0.01 compared to 0 nM.

#### Mitochondrial function in C2C12 myotubes directly treated with PCB126 or treated with the CM of PCB126-treated adipocytes

It was previously shown that inflammation plays a role in the development of muscle mitochondrial dysfunction (Cherry & Piantadosi, 2015; Remels et al., 2010, 2014), and it was demonstrated that PCB126 exposure in rats decreased muscle mitochondrial function (Tremblay-Laganière et al., 2019). To determine whether this decreased muscle mitochondrial function in rats exposed to PCB126 could be explained by the increased adipose tissue inflammation induced by the pollutant, we determined if exposure to CM from 3T3-L1 adipocytes exposed to PCB126 altered mitochondrial function in C2C12 myotubes (Figure 3C). As a control condition, we also exposed C2C12 myotubes directly to PCB126 for 24hrs (Figure 3D). In contrast to our initial hypothesis, treatment of C2C12 myotubes with CM from IS or IR PCB126-treated adipocytes did not alter mitochondrial function (Figure 3C). Unexpectedly, direct exposure to 100 nM PCB126 significantly increased resting OCR, proton leak and/or maximal mitochondrial capacity in control myotubes (IS control myotubes: *P*=0.0037 and 0.0302, for resting OCR and maximal capacity, respectively, IR myotubes: *P*=0.0555 and 0.0438, for resting and proton leak OCR, respectively, Figure 3D).

### Effect of direct or indirect PCB126 exposure on glucose uptake in IS and IR adipocytes and myotubes

#### Glucose uptake in IS and IR adipocytes directly treated with PCB126

PCB126 exposure has been associated with reduced glucose uptake in L6 myotubes (Mauger et al., 2016) and inhibition of the insulin signaling pathway in the muscle of rats (Wang et al., 2010). Moreover, adipose tissue inflammation is associated with decreased insulin sensitivity in skeletal muscle (Matsuda & Shimomura, 2013; Rains & Jain, 2011; Steinberg, 2007). Hence, we determined the effect of direct and indirect exposure to PCB126 on basal and insulin-stimulated glucose uptake in adipocytes and myotubes. Whereas adipocyte glucose uptake was significantly increased by insulin in IS conditions (Figure 4A), there was no significant increase in glucose uptake in response to insulin in IR adipocytes (Figure 4B), confirming that our IR conditions resulted in the development of insulin resistance in adipocytes. PCB126 exposure did not significantly alter basal or insulin-stimulated glucose uptake in IS adipocytes (Figure 4A), but a 100 nM PCB126 exposure significantly decreased glucose uptake in IR adipocytes, in basal and insulin-stimulated conditions (PCB126 effect, *P*=0.0305, Figure 4B). This decreased glucose uptake in IR adipocytes exposed to 100 nM PCB126 was not linked to any alteration in the phosphorylation of proteins of the insulin signaling pathway (p-IRS1, p-Akt, p-AS160; Figure S6A).

**Figure 4.**
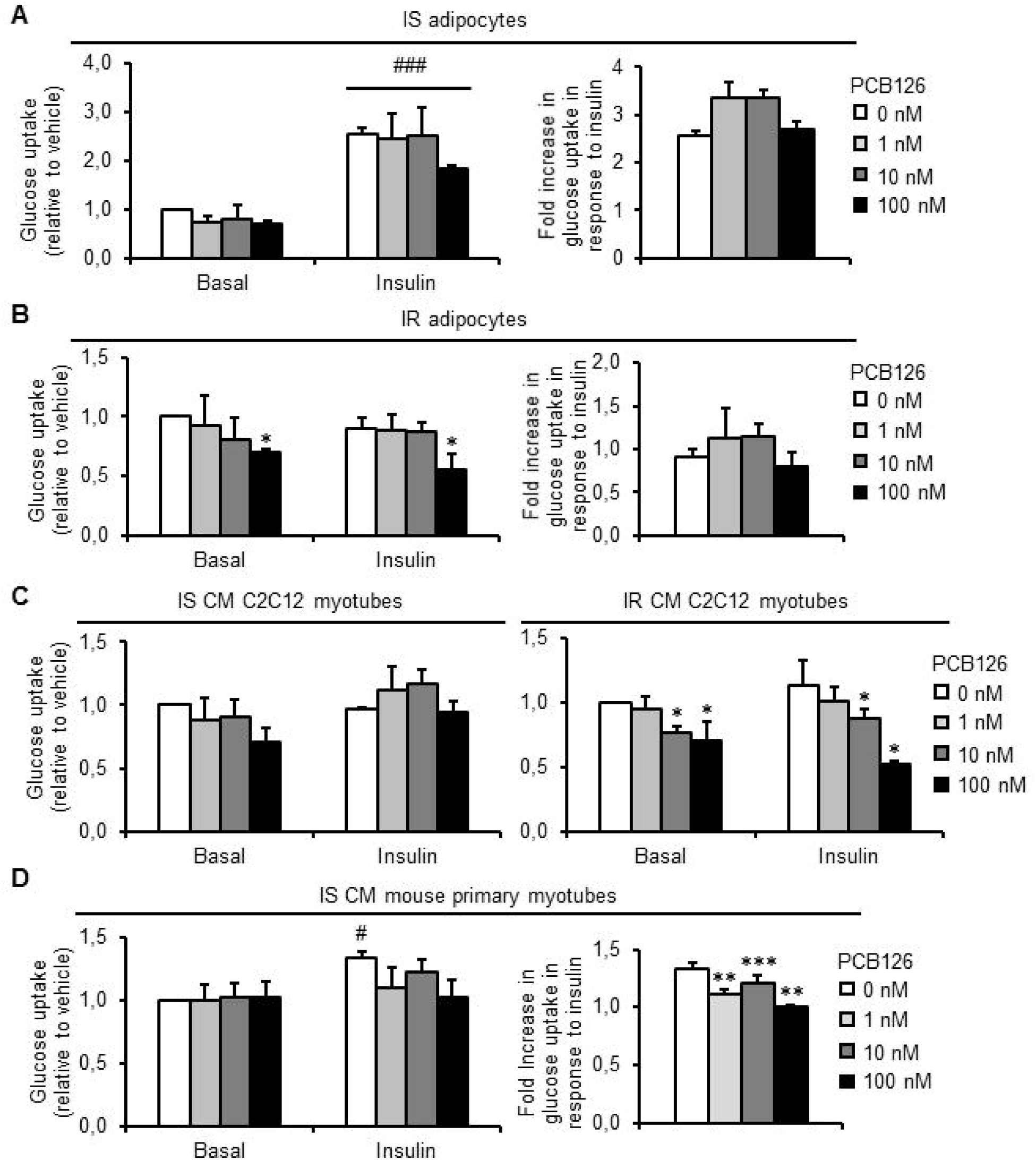
Glucose uptake in 3T3-L1 adipocytes exposed to PCB126 in different insulin sensitivity conditions and in C2C12 or mouse primary myotubes exposed to the conditioned medium (CM) of insulin sensitive (IS) and insulin resistant (IR) adipocytes treated with PCB126. A-B. 3T3-L1 adipocytes were differentiated in IS (A) and IR (B) conditions and treated for the last 24hrs of differentiation with different PCB126 concentrations. Left panel: glucose uptake, right panel: fold increase in glucose uptake in response to insulin. C. Differentiated C2C12 myotubes were exposed for the last 24hrs of differentiation to the CM of 3T3-L1 adipocytes exposed to different PCB126 concentrations in IS or IR conditions. D. Differentiated mouse primary myotubes were exposed for the last 24hrs of differentiation to the CM of 3T3-L1 adipocytes exposed to different PCB126 concentrations in IS conditions. Left panel: glucose uptake, right panel: fold increase in glucose uptake in response to insulin. A-D. After differentiation and treatments, cells were subsequently treated ±100 nM insulin for 20 min and exposed to 10 μM 2-deoxy-glucose and 0.5 μCi/mL [^3^H]2-deoxyglucose for 10 min. The absolute values of 2-deoxyglucose uptake under basal state (no insulin, no PCB126) were between 20 and 50 pmol/min/μg in 3T3-L1 adipocytes, between 50 and 80 pmol/min/μg in C2C12, and between 10 and 40 pmol/min/μg in mouse primary myotubes. Data are presented relative to the vehicle as mean ±SEM. n=3-4 independent experiments, each independent experiment was done in 3 replicates. *: *P*<0.05, **: *P*<0.01, ***: *P*<0.001 compared to 0 nM; #: *P*<0.05 compared with basal condition (no insulin).

#### Glucose uptake in C2C12 myotubes directly treated with PCB126 or treated with the CM of PCB126-treated adipocytes

We then wanted to determine whether exposure to CM from 3T3-L1 adipocytes exposed to PCB126 altered glucose uptake in C2C12 myotubes (Figure 4C). As a control condition, we also exposed C2C12 myotubes directly to PCB126 for 24hrs (Figure S5A). In control C2C12 myotubes, direct exposure to PCB126 did not significantly alter glucose uptake (Figure S5A). Exposure to CM from IS 3T3-L1 adipocytes (i.e. indirect PCB126 exposure) did not significantly alter glucose uptake in IS CM C2C12 myotubes, in either basal or insulin conditions (Figure 4C, left panel). In contrast, exposure to CM from IR 3T3-L1 adipocytes treated with 10 and 100 nM PCB126 significantly decreased glucose uptake in IR CM C2C12 myotubes, in basal and insulin conditions (*P*=0.0349 and *P*=0.0006, respectively) (Figure 4C, right panel). As in IR adipocytes treated with PCB126, the decreased glucose uptake in IR CM C2C12 myotubes was not linked to any alteration of the phosphorylation of proteins of the insulin signaling pathway (p-Akt, p-GSK3, Figure S6B).

#### Glucose uptake in mouse primary myotubes directly treated with PCB126 or treated with the CM of PCB126-treated adipocytes

It is recognized that insulin-stimulated glucose uptake is weak in C2C12 myotubes (Nedachi & Kanzaki, 2006), and as expected, insulin treatment did not significantly increase glucose uptake in this model (Figures 4C and S5A). Since one objective of the present study was to determine whether exposure to CM from PCB126-treated adipocytes altered insulin-stimulated glucose uptake in myotubes, we then used mouse primary myotubes, a muscle cell model more sensitive to insulin. Surprisingly, in IS control myotubes, direct exposure to 10 nM PCB126 tended to increase basal glucose uptake without reaching significance (Figure S5B, *P*=0.0749). In IR control myotubes, insulin did not increase glucose uptake suggesting that the high insulin levels in IR conditions also induced insulin resistance in mouse primary muscle cells (Figure S5B). As a consequence, the effect of CM from IR adipocytes treated with PCB126 on insulin-stimulated glucose uptake in mouse primary myotubes cannot be determined in IR conditions since the cells were already insulin resistant without any treatments. We then determined whether the CM from PCB126-treated adipocytes altered glucose uptake in mouse primary myotubes. A 24hr-exposure to the CM from IS or IR adipocytes did not significantly alter basal glucose uptake in mouse primary myotubes compared to myotubes treated with the CM of vehicle-treated adipocytes (i.e 0 nM PCB126) (Figure 4D and Figure S5C). However, the fold-increase in glucose uptake in response to insulin was decreased in mouse primary myotubes exposed to the CM of PCB126-treated IS 3T3-L1 adipocytes, suggesting insulin resistance development (1, 10 and 100 nM PCB126, *P*<0.01, Figure 4D).

### Effect of direct or indirect PCB126 exposure on glycolytic rate in IS and IR adipocytes and myotubes

#### Glycolysis in IS and IR adipocytes directly treated with PCB126

To further study the effect of 24hr-exposure to PCB126 on glucose metabolism, we next determined whether the impaired glucose uptake in IR adipocytes exposed to PCB126 could alter glycolytic rate (Figure 5). In accordance with glucose uptake results, a 24hr-exposure to PCB126 did not alter glycolytic rate in IS adipocytes (Figure 5A, left panel), but resting glycolytic rate and maximal glycolytic capacity were significantly decreased in IR adipocytes exposed to 1-100 nM PCB126 (for 1 nM *P*=0.0008 and 0.0007, for 10 nM *P*=0.0012 and 0.0018, and for 100 nM *P*=0.0029 and 0.0077, for resting and maximal glycolytic capacity, respectively) (Figure 5A, right panel).

**Figure 5.**
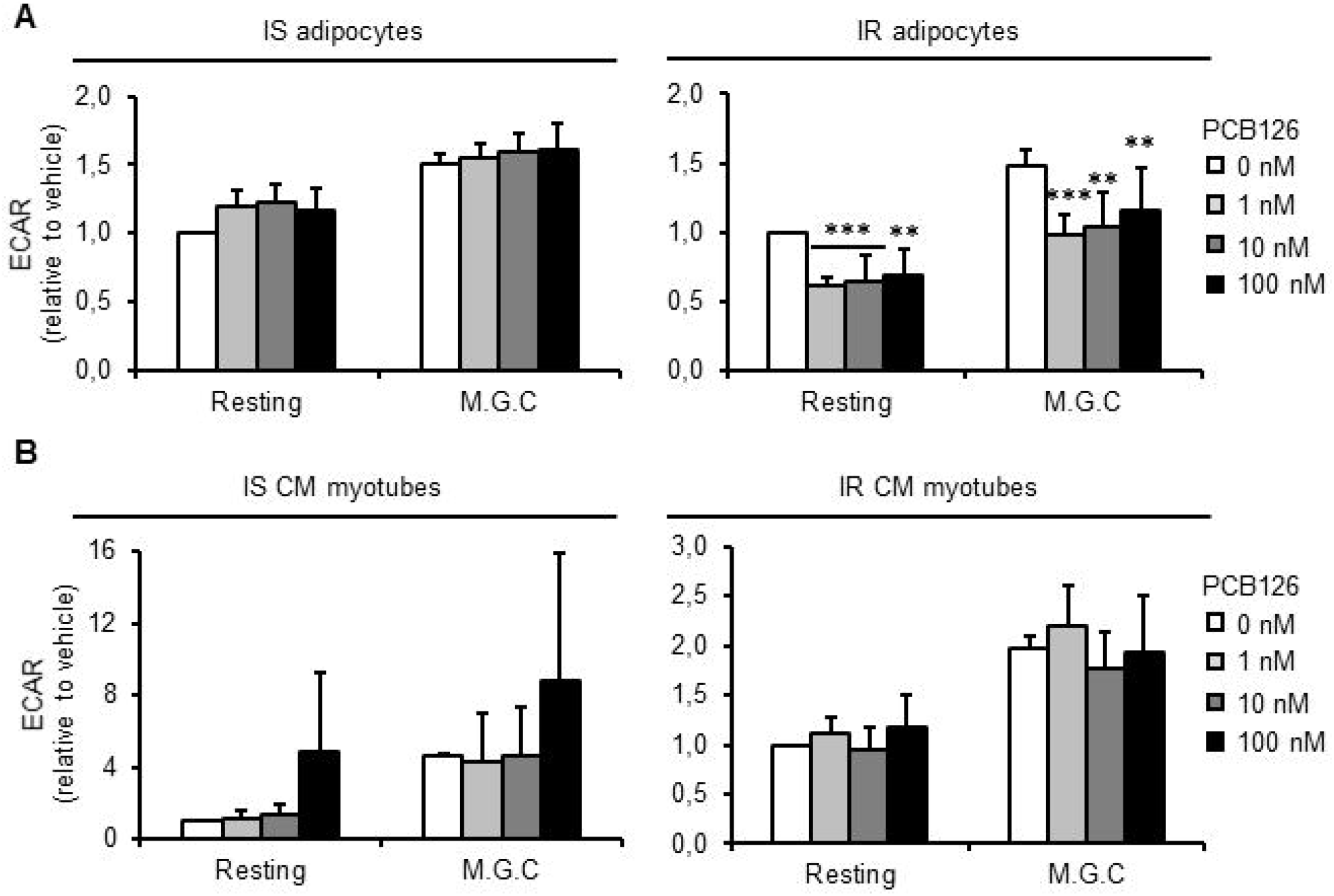
Glycolysis rates in 3T3-L1 adipocytes exposed to PCB126 in different insulin sensitivity conditions and in C2C12 exposed to the conditioned medium (CM) of insulin sensitive (IS) and insulin resistant (IR) adipocytes treated with PCB126. A. 3T3-L1 adipocytes were differentiated in IS (left panel) and IR (right panel) conditions and treated for the last 24hrs of differentiation with different PCB126 concentrations. B. Differentiated C2C12 myotubes were exposed for the last 24hrs of differentiation to the CM of 3T3-L1 adipocytes exposed to different PCB126 concentrations in IS (left panel) or IR conditions (right panel). A-B. Glycolysis rates were estimated by measuring extracellular acidification rates (ECAR) with a Seahorse analyzer (Agilent). ECAR were first measured in resting conditions, and cells were then treated with 600 ng/mL oligomycin to determine maximal glycolytic capacity (M.G.C.). Data are presented relative to the vehicle as mean ±SEM. n=4 independent experiments, each independent experiment was done in 5 replicates. *: *P*<0.05, **, *P*<0.01, ***: *P*<0.001 compared to 0 nM.

#### Glycolysis in C2C12 myotubes directly treated with PCB126 or treated with the CM of PCB126-treated adipocytes

Myotubes exposed to CM from IS or IR adipocytes did not show any alteration in resting or maximal glycolytic rates (Figure 5B). Glycolytic rates were also not altered by direct exposure to the pollutant (Figure S7).

### Effect of direct or indirect PCB126 exposure on oxidative stress markers in IR adipocytes and myotubes

#### Expression of anti-oxidant enzymes in IR adipocytes directly treated with PCB126

AhR activation and inflammation induced by coplanar PCBs may play a role in reactive oxygen species (ROS) production (Barouki, Coumoul, & Fernandez-Salguero, 2007; Hennig et al., 2002), which can in turn alter glucose uptake / insulin sensitivity and mitochondrial function (Bonnard et al., 2008; Pessler, Rudich, & Bashan, 2001; Potashnik, Bloch-Damti, Bashan, & Rudich, 2003). To better determine whether oxidative stress may explain altered metabolism induced by PCB126 in IR conditions, we measured the levels of oxidative stress markers in IR adipocytes after a 24hr-exposure to PCB126. PCB126 at 100 nM significantly increased the expression of SOD2 in IR adipocytes (*P*=0.0288), whereas other oxidative stress markers were not significantly altered by PCB exposure (Figure 6A).

**Figure 6.**
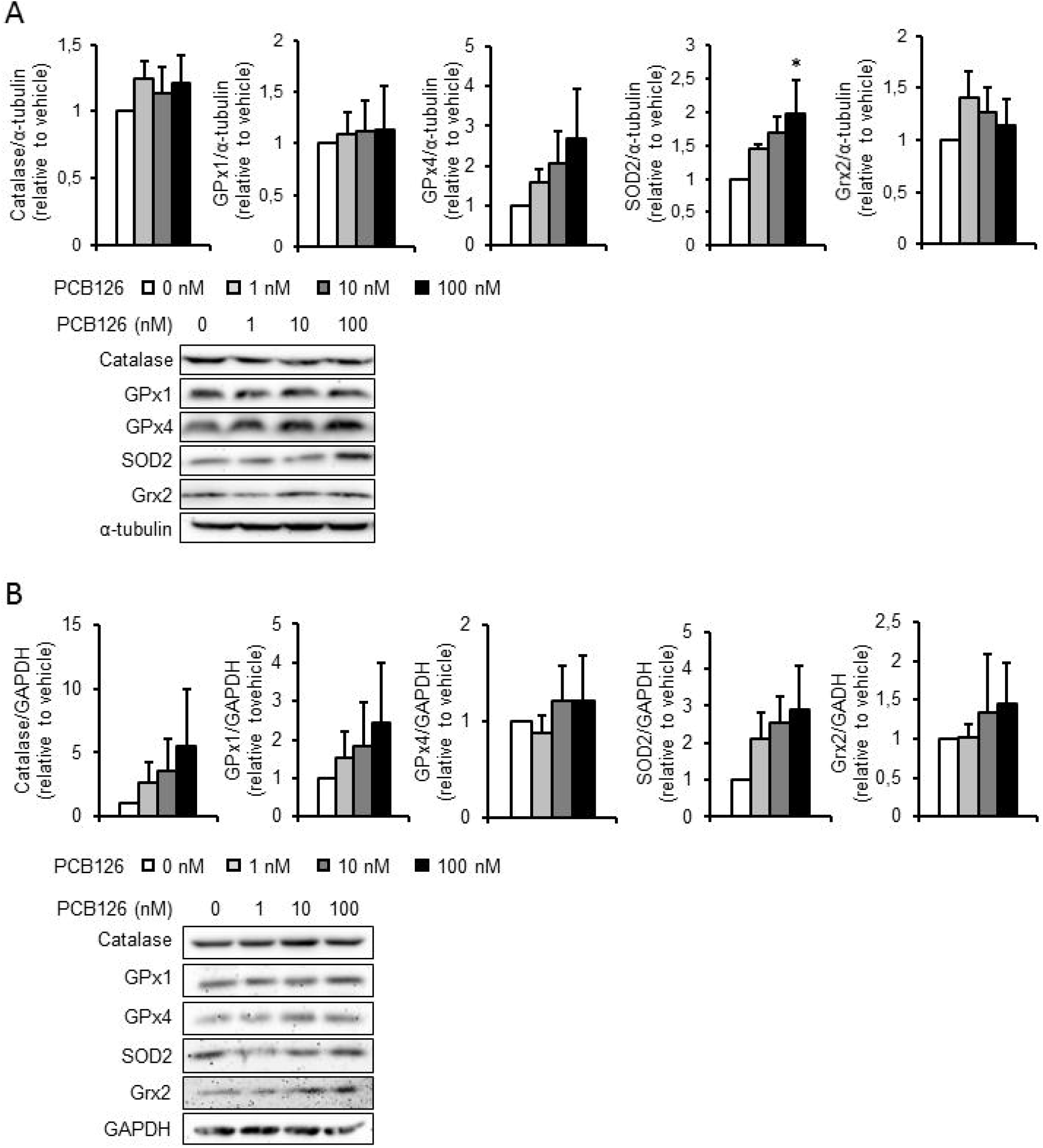
Oxidative stress markers in 3T3-L1 adipocytes exposed to PCB126 in insulin resistant (IR) conditions and in C2C12 myotubes exposed to the conditioned medium (CM) of PCB126-treated IR adipocytes. Levels of oxidative stress markers (catalase, glutathione peroxidase (GPx) 1 and 4, superoxide dismutase (SOD) 2, glutaredoxin (Grx) 2) in (A) 3T3-L1 adipocytes differentiated in IR conditions and exposed for 24hrs to different concentrations of PCB126 or (B) C2C12 myotubes exposed to the CM of PCB126-treated IR adipocytes. Top panel: quantification by density analysis, bottom panel: representative western blots. (A) α-tubulin and (B) GAPDH were used as loading controls. n=3-6 independent experiments. Data are presented relative to the vehicle as mean ±SEM. *: *P*<0.05 compared to 0 nM.

#### Oxidative stress markers in C2C12 myotubes treated with the CM of PCB126-treated IR adipocytes

In myotubes exposed to the CM of PCB126-treated IR adipocytes, the levels of catalase and SOD2 also tended to increase, but it did not reach significance (Figure 6B). The levels of other ROS detoxification enzymes were not altered in myotubes exposed to the CM of PCB126-treated IR adipocytes.

### Effect of direct or indirect PCB126 exposure on the active form of AMPK in IR adipocytes and myotubes

#### Expression of p-AMPK in IR adipocytes directly treated with PCB126

A key regulator of energy metabolism is AMPK. This kinase is known to be activated by an increased AMP/ATP ratio to restore energy status of the cell through the activation of ATP-producing pathways (e.g. glucose uptake, fatty acid oxidation, mitochondrial biogenesis) and the inhibition of ATP-consuming pathways (e.g. synthesis of fatty acids, glycogen, and amino acids) (Herzig & Shaw, 2018). To determine whether AMPK was involved in the measured decreased mitochondrial function and decreased glucose uptake upon exposure to PCB126 in IR adipocytes, we measured the levels of the activated form of AMPK (i.e. p-AMPK). IR adipocytes exposed to 100 nM PCB126 showed a significant decrease in the levels of p-AMPK (*P*=0.0078, Figure 7A).

**Figure 7.**
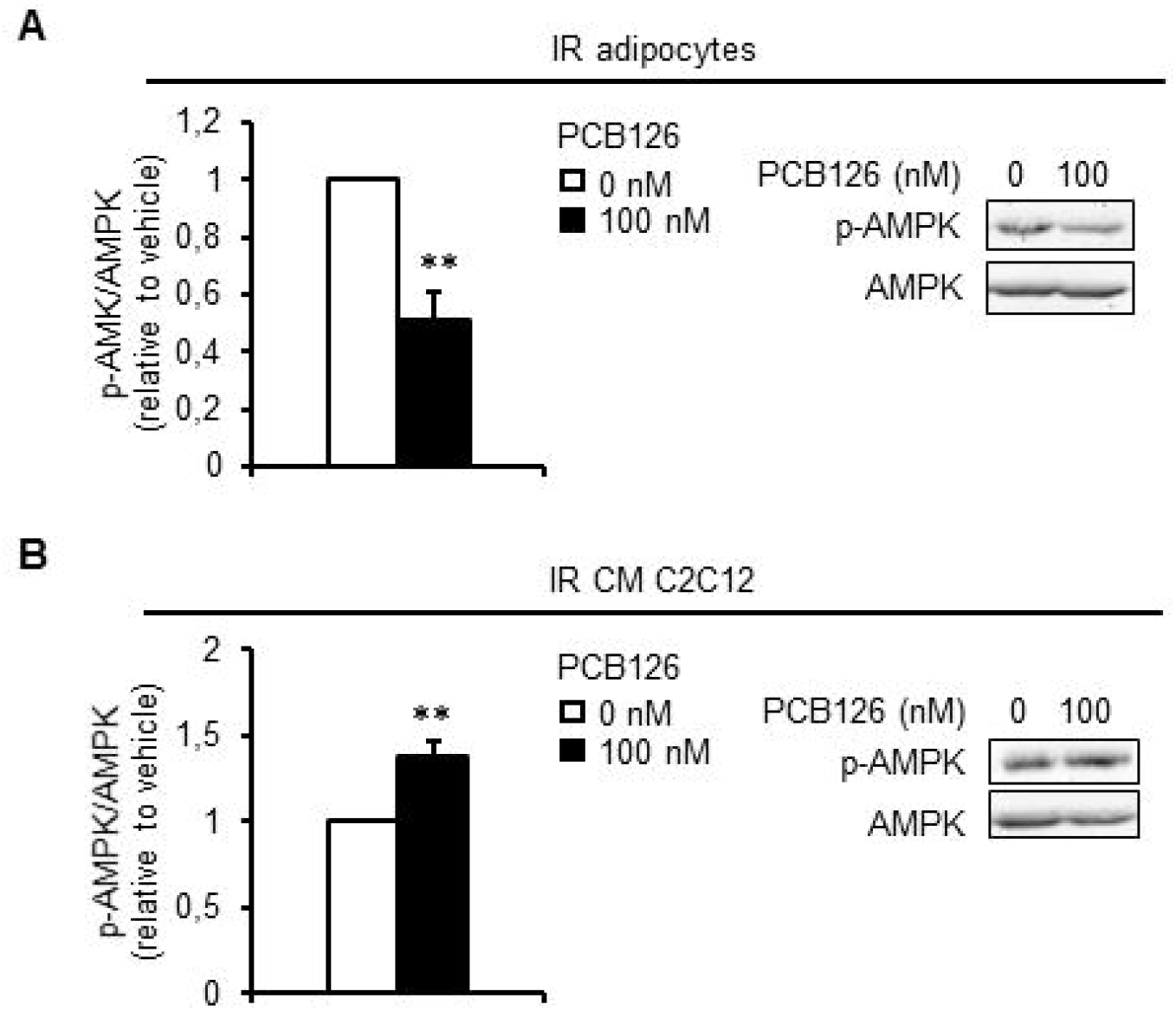
AMP-activated protein kinase (AMPK) levels in 3T3-L1 adipocytes exposed to PCB126 in insulin resistant (IR) conditions and in C2C12 myotubes exposed to the conditioned medium (CM) of PCB126-treated IR adipocytes. Levels of p-AMPK/AMPK in (A) 3T3-L1 adipocytes differentiated in IR conditions and exposed for 24hrs to 100 nM of PCB126 or (B) C2C12 exposed to the CM of IR adipocytes exposed to 100 nM PCB126. Left panel: quantification by density analysis, right panel: representative western blots. n=3 independent experiments. Data are presented relative to the vehicle as mean ±SEM. **: *P*≤0.01 compared to 0 nM.

#### Expression of p-AMPK in IR adipocytes directly treated with PCB126

We then determined whether the altered glucose uptake in C2C12 myotubes exposed to the CM of PCB126-treated IR adipocytes was also linked to a decreased AMPK activation. Curiously, in C2C12 myotubes exposed to the CM of PCB126-treated IR adipocytes, p-AMPK levels were significantly increased rather than decreased (*P*=0.011, Figure 7B)

## Discussion

Exposure to PCB126 has been associated with the development of insulin resistance and mitochondrial dysfunction (Connell, Singh, & Chu, 1999; Kim et al., 2017; Mauger et al., 2016; Tremblay-Laganière et al., 2019; Wakabayashi & Karbowski, 2001). PCB126 is a lipophilic compound that increases inflammation in adipose tissue (Gadupudi, Gourronc, Ludewig, Robertson, & Klingelhutz, 2015; Gourronc, Robertson, & Klingelhutz, 2017), and increased adipose tissue inflammation is one potential cause of insulin resistance and mitochondrial dysfunction development in skeletal muscle (Bhatnagar et al., 2010; Hwang et al., 2016). The general objective of the present study was to determine the role of adipose-to-muscle communication in the development of mitochondrial dysfunction and glucose metabolism alterations in skeletal muscle when exposed to PCB126. We showed that PCB126 altered the expression/secretion of adipokines and altered adipocyte mitochondrial function and glucose uptake, especially when adipocytes were rendered to be insulin resistant. These metabolic alterations were accompanied by increased SOD2 levels, known to be involved in reactive oxygen species detoxification, and decreased activation of AMPK, an important metabolic sensor. Furthermore, myotubes exposed to the CM of IR adipocytes treated with PCB126 showed altered glucose uptake compared to myotubes exposed directly to PCB126. However, exposure of myotubes to the CM of PCB126-treated adipocytes did not alter their mitochondrial function and showed an increased p-AMPK levels.

### Effect of pre-established insulin resistance on the metabolic response of adipocytes to PCB126 treatment

One objective of this study was to determine the effects of insulin resistance on the response of adipocytes to PCB126-exposure. Indeed, exposure to pollutants induces more significant metabolic defects in rodents developing obesity following a high-fat diet compared to their lean counterparts (Gray et al., 2013; Lim et al., 2009). For example, PCBs exacerbate hyperinsulinemia and insulin resistance in obese insulin resistant mice compared to lean insulin sensitive mice (Gray et al., 2013), suggesting increased sensitivity to pollutants when mice are already insulin resistant. Of note, in contrast to IS adipocytes, IR adipocytes exposed to PCB126 showed an increased secretion of cytokines/adipokines, including TNF-α, IL-6, leptin, adiponectin, and MCP-1. These results suggest that pre-established insulin resistance in adipocytes makes adipocytes more sensitive to the toxic effect of PCB126. This might be due to a better regulation of inflammatory response in metabolically healthy adipocytes. The present study focussed on the effect of insulin sensitivity and PCB126 on pro-inflammatory cytokines but how insulin resistance and/or PCB126 alter the secretion of anti-inflammatory cytokines such as IL-10 and IL-13 is also of interest and should be explored in future studies. Furthermore, it is important to note that, despite previous reports showing decreased adiponectin mRNA levels and protein secretion in adipocytes exposed to coplanar PCBs (Arsenescu, Arsenescu, King, Swanson, & Cassis, 2008; Gadupudi et al., 2015), in the present study, adiponectin secretion was increased in IR adipocytes exposed to PCB126. The differences between our study and others can be due to a difference in the adipocyte model used (3T3-L1 *vs*. human adipocytes), in the concentration of coplanar PCB used (30 to 100 times lower in the present study) or the timing of PCB treatment (in previous studies, adipocytes were treated with coplanar PCBs during the whole differentiation process, whereas in the present study, adipocytes were only treated for 24h, when the cells were already fully differentiated).

To the best of our knowledge, the effects of PCB126 exposure on glucose metabolism and mitochondrial function in adipocytes had never been studied. Therefore, the present study provides new insight into the role of PCB126 in adipose tissue metabolic dysfunction. The increased inflammation in IR adipocytes exposed to 100 nM PCB126 was associated with decreased glucose uptake. Mitochondrial function and glycolysis were also altered in IR adipocytes exposed to PCB126 at concentrations as low as 1 nM, which did not correspond to increased inflammation. Therefore, it seems that increased inflammation was probably not the cause of altered mitochondrial function and glycolysis in IR adipocytes. More studies are thus needed to better understand the mechanism by which PCB126 disrupts adipocyte energy metabolism. Importantly, exposure to PCB126 did not alter these metabolic pathways in IS adipocytes, showing that insulin sensitivity status might influence the response of adipocytes to PCB126 exposure.

The altered glucose metabolism and mitochondrial function in PCB126-treated IR adipocytes were associated with increased adipokine secretion and increased level of SOD2. Interestingly, increased inflammation and oxidative stress are recognized to promote mitochondrial dysfunction and to alter glucose metabolism in adipocytes (Fazakerley et al., 2018; Manna & Jain, 2015; Paglialunga, Ludzki, Root-McCaig, & Holloway, 2015; Steinberg, 2007). In the present study, oxidative stress has been estimated by measuring the levels of ROS detoxification enzymes. To better study the role of PCB126 in oxidative stress development in adipocytes, future studies should explore the impact of PCB126 on other important players in the modulation of oxidative stress such as glutathione levels.

The negative effect of PCB126 on glucose metabolism and mitochondrial function in IR adipocytes was also associated with decreased activation of AMPK (i.e. its phosphorylation levels). AMPK is an important metabolic sensor known to activate glucose uptake independently of the insulin signaling pathway and to increase mitochondrial activity (Crawford et al., 2010; Jäer, Handschin, St-Pierre, & Spiegelman, 2007). Previously, a link between altered AMPK activation and increased inflammation of adipose tissue has been demonstrated (Jung, Park, Choi, Kim, & Lee, 2018; Yang et al., 2016; Zhao et al., 2018). Our results therefore suggest that the decrease in glucose uptake and mitochondrial function in IR adipocytes exposed to 100 nM PCB126 might be the result of an alteration of the AMPK pathway because of increased inflammation / oxidative stress.

### Role of adipocyte-secreted factors in the development of muscle cell mitochondrial dysfunction when exposed to PCB126

Despite the role of skeletal muscle in the maintenance of glucose homeostasis, the effect of PCB126 or other environmental pollutants on skeletal muscle energy metabolism has not been well studied. High circulating levels of pollutants have been associated with decreased mitochondrial enzyme activity in human skeletal muscle (Imbeault et al., 2002). Furthermore, our team showed that PCB126 exposure in rats was associated with decreased mitochondrial function in muscle (Tremblay-Laganière et al., 2019), which was not reproduced when muscle cells were directly exposed to the pollutant (Mauger et al., 2016). Here, we tested whether the adipose-to-muscle communication in the context of PCB126 exposure could induce mitochondrial dysfunction in skeletal muscle. However, CM from PCB126-treated IS or IR adipocytes did not alter mitochondrial function in C2C12 myotubes. Therefore, with the present model, we cannot confirm that the adipose-to-muscle communication is responsible for the decreased mitochondrial function in muscle from rats exposed to PCB126. Curiously, the activation of AMPK (i.e. p-AMPK levels) was increased rather than decreased in C2C12 myotubes exposed to the CM of IR-treated adipocytes. Since AMPK is known to activate mitochondrial function, it is possible that C2C12 myotubes were able to maintain their mitochondrial function upon CM exposure because of increased AMPK activity.

### Direct effect of PCB126 on glucose metabolism in muscle cells

We previously showed decreased glycolytic function and glucose uptake in L6 myotubes exposed for 24hrs to PCB126 (Mauger et al., 2016). Moreover, exposure to a PCB mixture (Aroclor 1254) altered skeletal muscle insulin signaling pathway and GLUT4 translocation in rats (Williams et al., 2013). In the present study, C2C12 or mouse primary myotubes directly exposed to the same concentrations of PCB126 than in our previous study using L6 myotubes (Mauger et al., 2016) showed no alteration of glucose uptake and glycolytic rates. These different effects of PCB126 on muscle cell glucose metabolism might be due to metabolic differences between L6, C2C12 and mouse primary myotubes. It is known that C2C12 lack insulin-responsive GLUT4 vesicles required for significant insulin-stimulated glucose uptake (Tortorella & Pilch, 2002), while L6 and primary mouse muscle cells possess these vesicles (Robinson, Robinson, James, & Lawrence, 1993; Tortorella & Pilch, 2002). In addition, under conditions similar to ours (1 g/L glucose and 2% FBS), C2C12 myotubes express a greater proportion of slow myosin heavy chains (Artaza et al., 2002; J B Miller & Stockdale, 1986; Jeffrey Boone Miller, 1990). Thus, C2C12 myotubes may have a more oxidative metabolism than L6 myotubes. This hypothesis has been confirmed in a recent study in which it was demonstrated that L6 cells have lower mitochondrial and fatty acid oxidation capacities than C2C12 cells, while having higher emission of ROS (Robinson et al., 2019). Future studies should therefore determine whether the higher glycolytic capacity and higher ROS emission in L6 cells may explain the fact that this muscle cell model is more sensitive to PCB126 than C2C12 when exposed to PCB126.

### Role of adipocyte-secreted factors in the development of altered muscle cell glucose uptake when exposed to PCB126

Using the CM from PCB126-treated IR adipocytes, we showed that the adipocyte secretome decreased basal glucose uptake in C2C12 myotubes, whereas the CM from PCB126-treated IS adipocytes altered insulin sensitivity in mouse primary myotubes. Even if this altered glucose transport in myotubes was not associated with any alteration of the insulin signaling pathway, it is possible that GLUT4 content/translocation and/or GLUT1 levels/activity were affected. Further studies are thus needed to determine whether this altered glucose uptake is the result of an alteration of the expression or activity of the glucose transporters GLUT1 and/or GLUT4.

Previous studies have demonstrated that lipids can have a detrimental effect on muscle metabolism and may even induce insulin resistance (Aguer et al., 2015; Coles, 2016). However, in our model, the altered glucose uptake in myotubes exposed to the CM of PCB126-treated IS and IR adipocytes was probably not due to FFA since lipolysis rate was decreased rather than increased in IS adipocytes exposed to PCB126, and lipolysis rate was not altered in IR adipocytes exposed to PCB126.

Adipose tissue inflammation may be another player in the development of altered muscle glucose transport. Adipocytes secrete adipokines involved in autocrine/paracrine and endocrine functions. Adipokines alter metabolic responses locally in adipose tissue, as well as in distant tissues such as skeletal muscle (Coles, 2016; Luo & Liu, 2016; Scherer, 2006). Under healthy conditions, adipokines maintain energy homeostasis, but dysregulation of adipokine secretion causes lipotoxicity in skeletal muscle (Coles, 2016). In this sense, adipose tissue dysfunction has been associated with increased inflammation, which may explain the association between obesity and the increased risk of developing insulin resistance and type 2 diabetes. Of note, in the present study, PCB126 promoted inflammation in IR adipocytes in association with decreased basal glucose uptake in myotubes exposed to IR adipocyte secretome. Similar results were obtained when muscle cells were treated directly with different adipocytokines, such as TNF-α, MCP-1 and IL-6 (Li et al., 2017; Sell, Dietze-Schroeder, Kaiser, & Eckel, 2006). Taken together, the results from the present study and others suggest that PCB126-induced inflammation in adipose tissue might be responsible for the decreased insulin response and reduced basal glucose uptake in muscle cells when exposed to PCB126 (Gadupudi et al., 2015; Gourronc et al., 2017; Williams et al., 2013). Interestingly, treatment with CM from hypoxia-treated 3T3-L1 adipocytes also induced insulin resistance in C2C12 myotubes (Yu et al., 2011), suggesting that different stressors might impact adipokine secretion, which in turn alter muscle insulin sensitivity. On the other hand, PCB126 treatment in IS adipocytes altered mRNA adiponectin expression but did not affect the secretion of adiponectin or other adipokines, suggesting that the effect of IS CM on myotube insulin sensitivity might be due to other secreted factors than the ones we measured in the present study. Further research is thus needed to determine which factor(s) secreted by PCB126-exposed IS adipocytes affect myotube insulin sensitivity. The potential candidates include adipocytokines not measured in our study (e.g. IL-1, IL-8, keratinocyte chemoattractant-1), metabolites that could be differentially secreted by adipocytes when exposed to PCB126, as well as miRNAs.

### Limitations

Our conclusions are limited by our model that studied a one-way communication between adipocytes and myotubes using cell cultures *in vitro*. We acknowledge that this does not fully represent what happens in an organism where the crosstalk between muscle and adipose tissue may also regulate metabolism (Bogdanowicz & Lu, 2013; Li et al., 2017; Pandurangan, Jeong, Amna, Van Ba, & Hwang, 2012). Our model was appropriate for our objectives in determining the role of PCB126 on adipokine secretion from adipocytes and how this alters muscle energy metabolism. However, it needs to be considered as an isolated system that possesses some limitations. Indeed, at the whole-body level, PCBs will also alter the secretion of cytokines and the production of ROS from other tissues and cell types, such as the liver, immune cells and endothelial cells (Cocco et al., 2015; Lim et al., 2009; Tremblay-Laganière et al., 2019), which can also alter muscle metabolism. Moreover, *in vivo*, PCBs might be metabolized by the liver, which is not taken into account in our model. Lastly, our study focussed on a PCB126 concentration (100 nM) that is higher than estimated environmental exposures, which may limit the generalization of our results. However, we studied the acute effect of a single pollutant (i.e. exposure for 24hrs to PCB126), while humans are chronically exposed to several pollutants that may act in a similar fashion to PCB126. Further studies are thus required to determine whether exposure to a mix of pollutants for a longer period shows similar effects on adipose tissue metabolism and on adipose-to-muscle communication.

## Conclusion

In summary, we demonstrated that PCB126 promotes inflammation and metabolic defects in adipocytes in relation with decreased AMPK activity, particularly when those cells were already insulin resistant before exposing them to the pollutant. Moreover, our data suggest that the alteration of glucose uptake and insulin response in skeletal muscle in response to PCB126 treatment is the result of PCB126-induced inflammation in adipose tissue. More broadly, our study shows the importance of the cross-talk between adipose tissue and skeletal muscle in the development of insulin resistance.

## Acknowledgment

The authors would like to thank Dr. Marc Foretz for his generous gift of mouse primary myoblasts, Dr. Mary-Ellen Harper for the use of the XF-96 analyzer (Seahorse Bioscience) and the provision of her lab for radiation experiments, and Marc Rigden for *Cyp1a1* mRNA determination. This work was funded by a Natural Sciences and Engineering Research Council of Canada (NSERC) Discovery Grant (2015–06263) to CA. Master scholarships to AC were provided by NSERC and Fonds de recherche du Québec – Santé and to LG by Institut du Savoir Montfort.

## Author contributions

Experiments were performed in Dr. Aguer’s laboratory at the Institut du Savoir Monfort (Ottawa, ON, Canada) and in Dr. Atlas’ laboratory at Health Canada (Ottawa, ON, Canada) except for Seahorse and radiation experiments that were conducted in Dr. Harper’s laboratory (Biochemistry, Microbiology, and Immunology Department, University of Ottawa, Canada). Conception and design of the experiments were done by CA; collection, assembly, analysis, and interpretation of data by AC, FA, VP, LG, EA, and CA; drafting the article or revising it critically for important intellectual content by AC, FA, VP, LG, EA, and CA, and approval of the final version by AC, FA, VP, LG, EA, and CA.

