## Supplemental figures for "PCB126-mediated effects on adipocyte energy metabolism and adipokine secretion may result in abnormal glucose uptake in muscle cells"

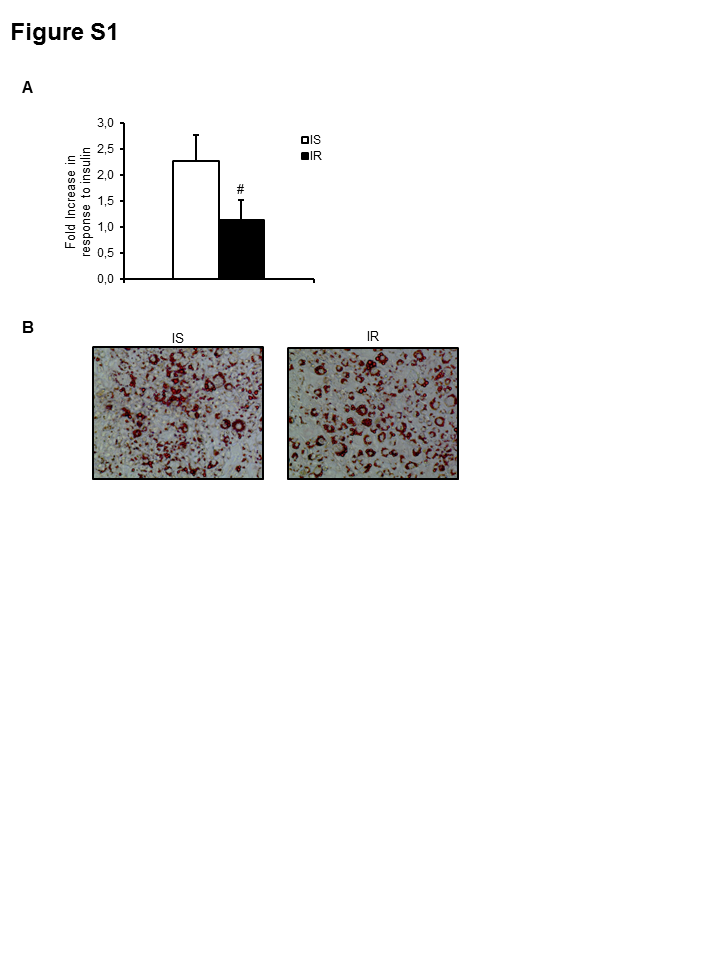

**Figure S1. Effect of insulin sensitive (IS) and insulin resistant (IR) conditions on 3T3-L1 adipocyte glucose uptake and lipid content.** A. Fold-increase in glucose uptake in response to insulin stimulation. Glucose uptake was measured in 3T3-L1 adipocytes differentiated in IS and IR conditions, under basal condition (no insulin) and after 20 min insulin treatment (100 nM). (#: *P*<0.05 compared to IS). B. Lipid droplet visualization by Oil Red O staining in 3T3-L1 adipocytes differentiated in IS or IR conditions. Lipid droplets were visualized using a light microscope with a 40X objective.

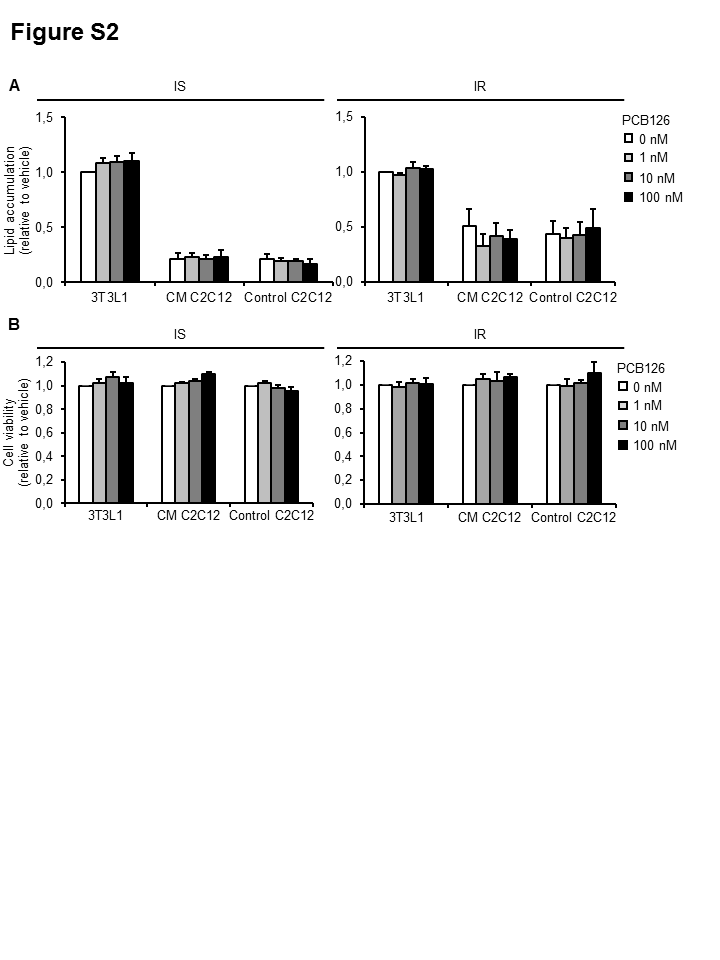

**Figure S2. Effect of PCB126 or conditioned medium (CM) treatments on lipid accumulation and cell viability in 3T3-L1 adipocytes and C2C12 myotubes.** A-B. Insulin sensitive (IS), left panel) and insulin resistant (IR, right panel) 3T3-L1 adipocytes and C2C12 myotubes (control C2C12) were directly exposed to different concentrations of PCB126 for 24hrs. The conditioned media from adipocytes was subsequently transferred to C2C12 myotubes for 24hrs (CM C2C12). A. After treatments, cells were fixed, and lipid droplets stained with Oil Red O. Stained lipid droplets were extracted with isopropanol and a triplicate for every well was read at 492 nm. Mean ±SEM. n=4 independent experiments, each independent experiment was done at least in triplicate. B. After treatment, cells were incubated with PrestoBlue for 30 min. Plates were read at 570 nm and 600 nm (reference wavelength). n=3 independent experiments, each independent experiment was done at least in triplicate. A. Data are presented relative to the vehicle (0 nM PCB126) in 3T3-L1. B. Data are presented relative to the vehicle (0 nM PCB126) for each cell line independently. Mean ±SEM.

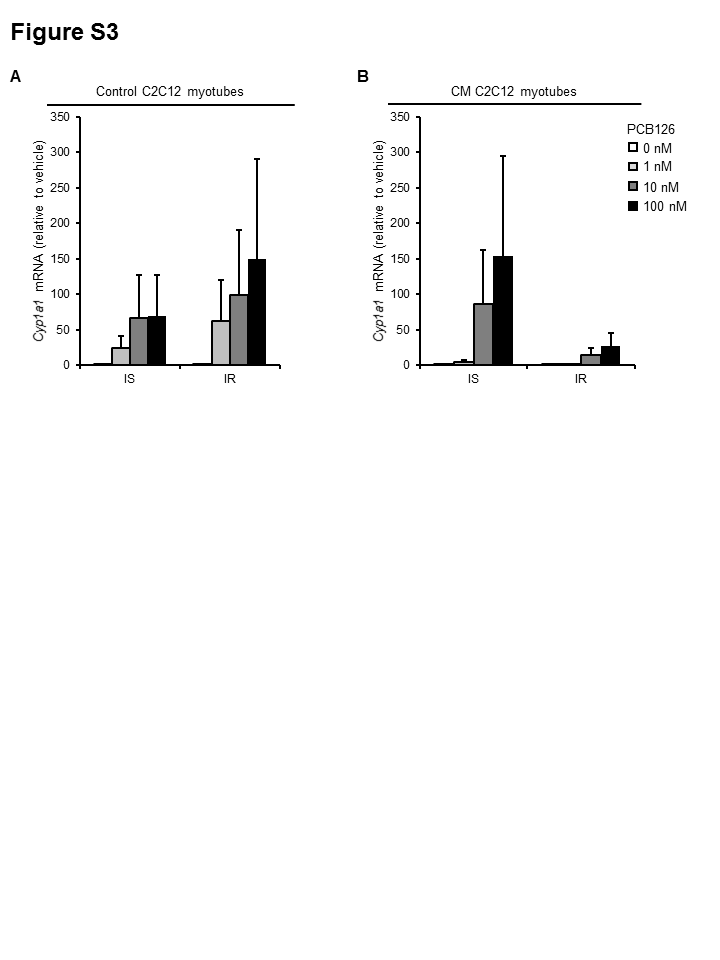

**Figure S3. Effect of direct or indirect PCB126 exposure on *Cyp1a1* mRNA expression in C2C12 myotubes.** Differentiated C2C12 myotubes were exposed for the last 24hrs of differentiation (A) to different PCB126 concentrations in insulin sensitive (IS) or insulin resistant (IR) conditions or (B) to the conditioned medium (CM) of 3T3-L1 adipocytes exposed to different PCB126 concentrations in IS or IR conditions. *Cyp1a1* mRNA levels were normalized to β-actin mRNA levels and analyzed using the ΔΔCT method. Average of normalized ΔΔCT is presented relative to the vehicle (IS, no PCB) ±SEM. (n=3 independent experiments, each independent experiment was done in triplicate).

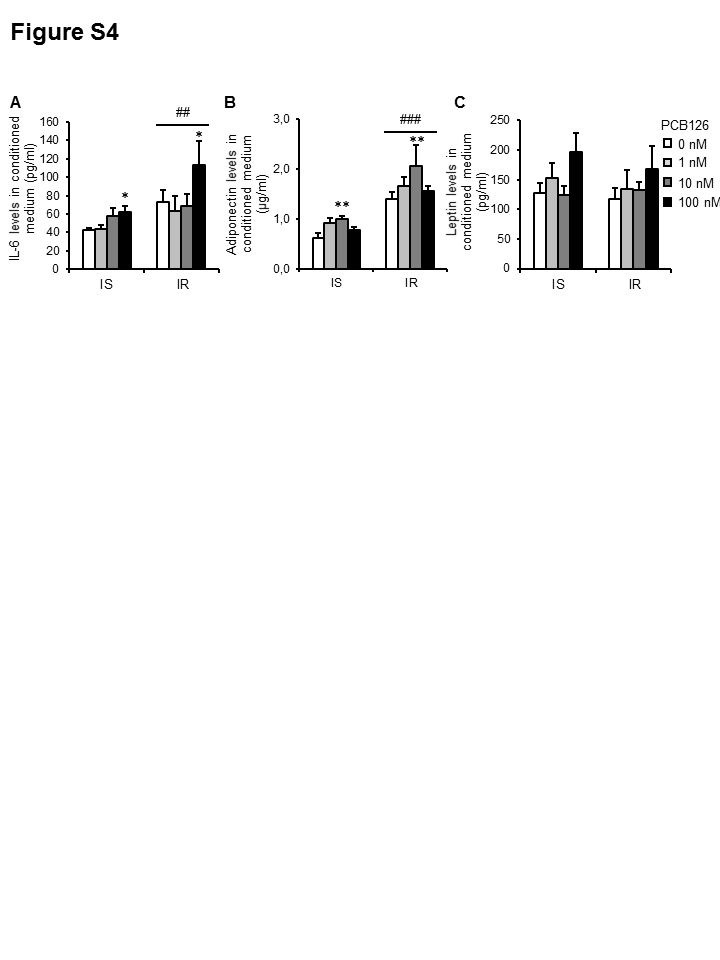

**Figure S4. Effect of PCB126 exposure and insulin resistance on adipokine secretion in 3T3-L1 adipocytes.** A-C. Insulin sensitive (IS) and insulin resistant (IR) 3T3-L1 adipocytes were exposed to different concentrations of PCB126 for 24hrs. Interleukin 6 (IL-6) (A), adiponectin (B) and leptin (C) levels were then measured in the conditioned medium using ELISA kits. Mean ±SEM. n=3-4 independent experiments, each independent experiment was done in duplicate. *: *P*<0.05, **: *P*<0.01 compared to vehicle-treated adipocytes, ##: *P*<0.01, ###: *P*<0.001, main effect of IR conditions.

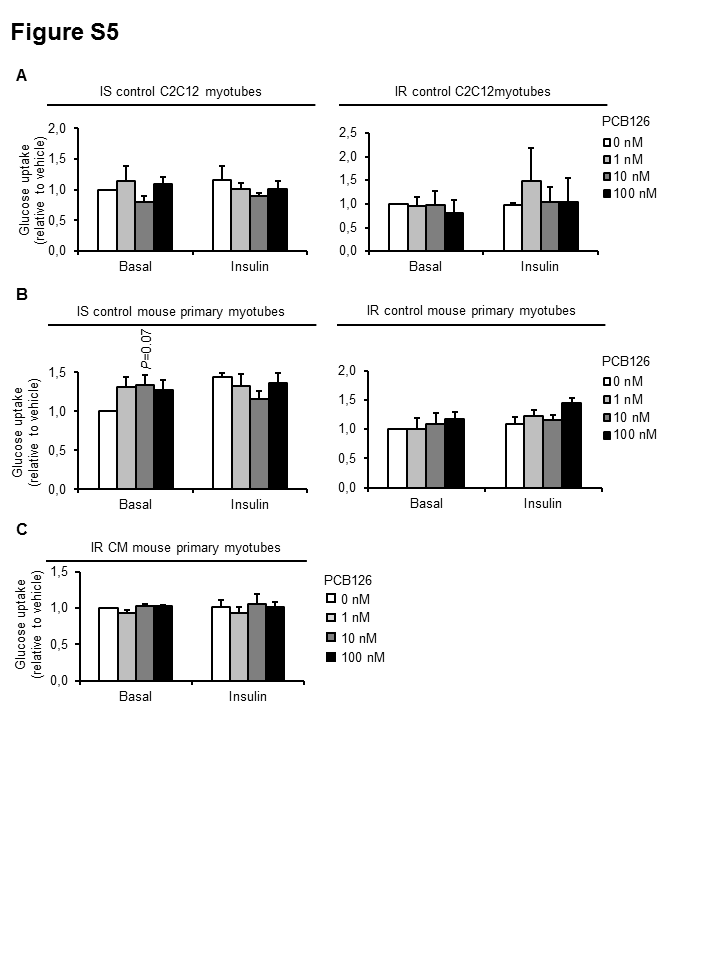

**Figure S5. Glucose uptake in C2C12 or mouse primary myotubes directly exposed to PCB126 in insulin sensitive (IS) and insulin resistant (IR) conditions.** Differentiated C2C12 myotubes (A) or mouse primary myotubes (B,C) were exposed for the last 24hrs of differentiation to different PCB126 concentrations in IS or IR conditions (A and B) or to the CM of PCB126-treated IR adipocytes for the last 24hrs of differentiation (C). After differentiation and treatments, cells were subsequently treated with ±100 nM insulin for 20 min and exposed to 10 μM 2-deoxy-glucose and 0.5 μCi/mL [^3^H]2-deoxyglucose for 10 min. Data are presented relative to the vehicle as mean ±SEM. n=3-4 independent experiments, each independent experiment was done in 3 replicates.

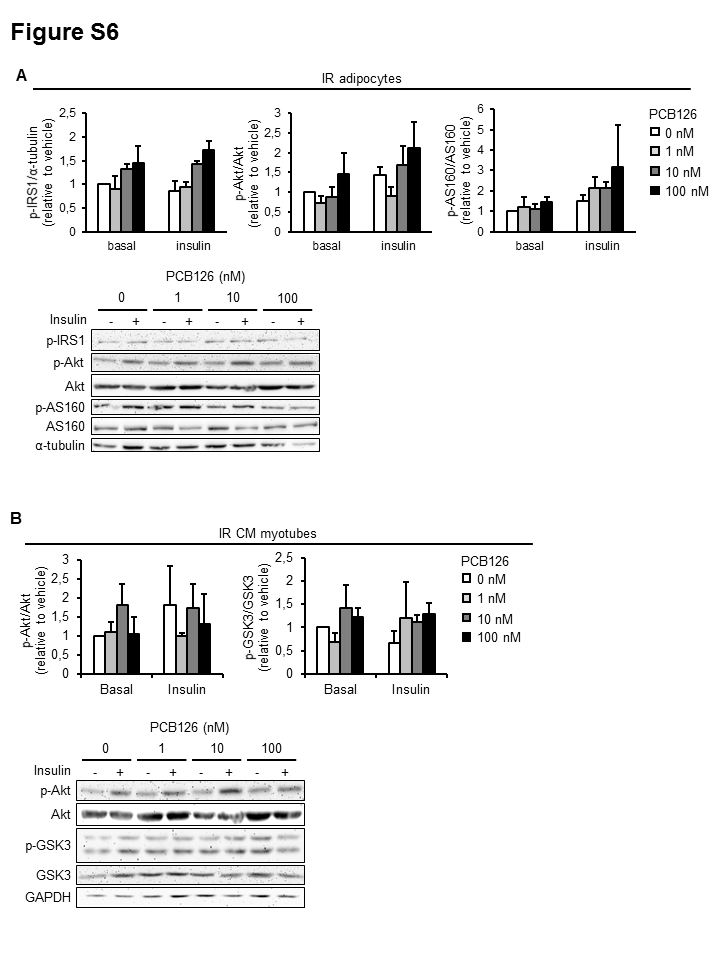

**Figure S6. Insulin signaling in 3T3-L1 adipocytes exposed to PCB126 in insulin resistant (IR) conditions and in C2C12 exposed to the conditioned medium (CM) of PCB126-treated IR adipocytes.** A. 3T3-L1 adipocytes were differentiated in IR conditions and treated for the last 24hrs of differentiation with different PCB126 concentrations and cells were subsequently treated with ±100 nM insulin. Top panel: quantification of p-IRS1/α-tubulin, p-Akt/Akt and p-AS160/AS160 by density analysis. Bottom panel: representative western blots. Tubulin was used as the loading control. B. Differentiated C2C12 myotubes were exposed for the last 24hrs of differentiation to the CM of PCB126-treated IR adipocytes and cells were subsequently treated with ±100 nM insulin. Top panel: quantification of p-Akt/Akt and p-GSK3/GSK3 by density analysis. Bottom panel: representative western blots. GAPDH was used as the loading control. Data are presented relative to the vehicle as mean ±SEM. n=3 independent experiments.

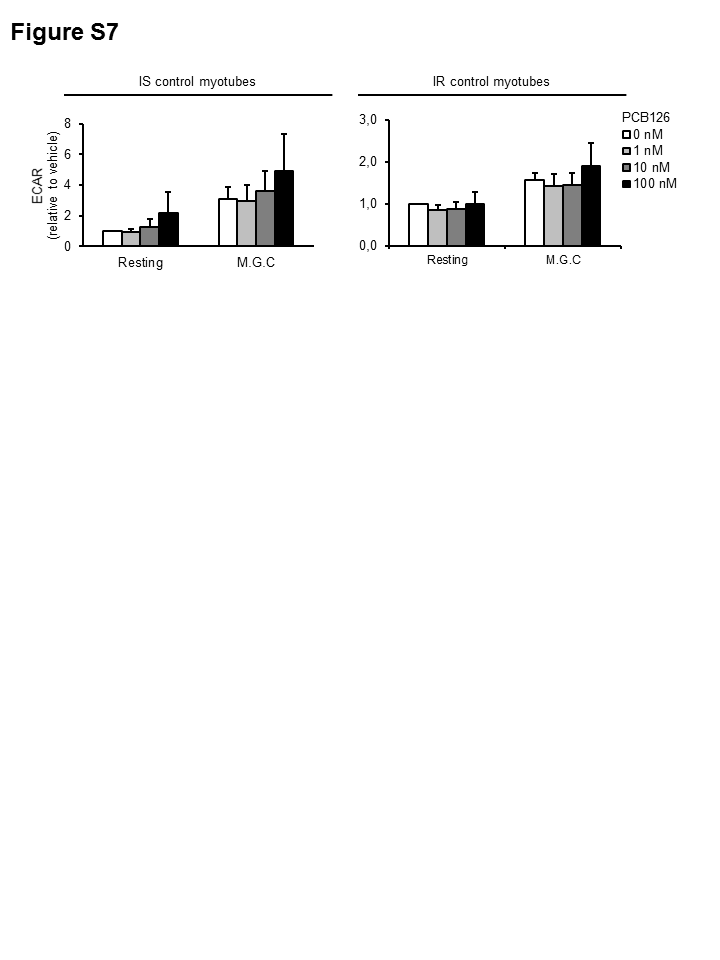

**Figure S7. Glycolytic rates in C2C12 myotubes directly exposed to PCB126 in insulin sensitive (IS) or insulin resistant (IR) conditions.** Differentiated C2C12 myotubes were exposed for the last 24hrs of differentiation to different PCB126 concentrations in IS (left panel) or IR conditions (right panel). Glycolytic rates were estimated by measuring extracellular acidification rates (ECAR) with a Seahorse analyzer (Agilent). ECAR were first measured in resting conditions, and cells were then treated with 600 ng/mL oligomycin to determine maximal glycolytic capacity (M.G.C.). Data are presented relative to the vehicle as mean ±SEM. n=4 independent experiments, each independent experiment was done in 5 replicates.
